# Paradoxical tumor suppressive role of the *musculoaponeurotic fibrosarcoma* gene in colorectal cancer

**DOI:** 10.1101/2022.08.25.505236

**Authors:** Hiroaki Itakura, Tsuyoshi Hata, Daisuke Okuzaki, Koki Takeda, Kenji Iso, Yamin Qian, Yoshihiro Morimoto, Tomohiro Adachi, Haruka Hirose, Yuhki Yokoyama, Takayuki Ogino, Norikatsu Miyoshi, Hidekazu Takahashi, Mamoru Uemura, Tsunekazu Mizushima, Takao Hinoi, Masaki Mori, Yuichiro Doki, Hidetoshi Eguchi, Hirofumi Yamamoto

**Affiliations:** Department of Surgery, Gastroenterological Surgery, Graduate School of Medicine, Osaka University, Yamadaoka 2-2, Suita, Osaka, 565-0871, JAPAN; Genome Information Research Centre, Research Institute for Microbial Diseases, Osaka University, Yamadaoka 3-1, Suita, Osaka, 565-0871, JAPAN; Laboratory of Human Immunology (Single Cell Genomics), WPI Immunology Research Center, Osaka University, Yamadaoka 3-1, Suita, Osaka, 565-0871, JAPAN; Department of Molecular Pathology, Division of Health Sciences, Graduate School of Medicine, Osaka University, Yamadaoka 1-7, Suita, Osaka, 565-0871, JAPAN; Department of Surgery, Hiroshima City North Medical Center Asa Citizens Hospital, 1-2-1, Kameyama-minami, Asakita-ku, Horoshima, 731-0293, JAPAN; Department of Surgery, Osaka Police Hospital, 10-31, Kitayama-town, Tennoji-ku, Osaka city, Osaka, 543-0035, JAPAN; Department of Clinical and Molecular Genetics, Hiroshima University Hospital, 1-2-3, Kasumi, Minami-ku, Hiroshima, 734-8551, JAPAN; Department of Surgery, Graduate School of Medical Sciences, Tokai University, 143, Shimokasuya, Isehara, Kanagawa, 259-1193, JAPAN

**Keywords:** Colorectal cancer, c-MAF, p53, carcinogenesis

## Abstract

Somatic cell reprogramming using the microRNAs miR200c, miR-302s, and miR-369s leads to increased expression of cyclin-dependent kinase inhibitors in human colorectal cancer (CRC) cells and suppressed tumor growth. Here, we investigated whether these microRNAs inhibit colorectal tumorigenesis in *CPC;Apc* mice, which are prone to colon and rectal polyps. Repeated administration of microRNAs inhibited polyp formation. Microarray analysis indicated that c-MAF, which reportedly shows oncogene-like behavior in multiple myeloma and T-cell lymphoma, decreased in tumor samples but increased in microRNA-treated normal mucosa. Immunohistochemistry identified downregulation of c-MAF as an early tumorigenesis event in CRC, with low c-MAF expression associated with poor prognosis. Of note, c-MAF expression and p53 protein levels were inversely correlated in CRC samples. *c-MAF* knockout led to enhanced tumor formation in azoxymethane/dextran sodium sulfate–treated mice, with activation of cancer-promoting genes. c-MAF may play a tumor-suppressive role in CRC development.

## Introduction

Colorectal cancer (CRC) is among the most prevalent cancers worldwide, ranking as the third most common cancer and the fourth leading cause of cancer death (Abbasi et al., 2015; Brenner et al., 2014; Torre et al., 2015). Despite considerable progress in surgery and chemotherapy in the past decade and the development of molecular targeted therapies, 5-year survival remains at 50% to 65% (Biller and Schrag, 2021; Colvin et al., 2014; Dienstmann et al., 2017; Torre et al., 2015).

Cancer is a genetic disease, but epigenetic alterations also are involved in its initiation and progression. By introducing Yamanaka reprogramming factors, *i.e.,* Oct3/4, Sox2, Klf4, and c-Myc, for generation of induced pluripotent stem cells (Takahashi and Yamanaka, 2006), we previously showed that reprogramming of CRC cells reduces their malignant potential (Miyoshi et al., 2010). In this way, reprogramming could prove useful as a cancer therapy. Other work has highlighted potential risks with virus vectors and the oncogenic *c-myc* gene (Jia et al., 2010; Kaji et al., 2009; Okita et al., 2008; Takahashi and Yamanaka, 2006; Woltjen et al., 2009). To sidestep these risks, we have used the microRNAs (miRNAs) miR-200c, miR-302s, and miR-369s to reprogram differentiated human and mouse somatic cells (Miyoshi et al., 2011). Our results showed that these miRNAs trigger increased expression of cyclin-dependent kinase inhibitors such as p16^Ink4a^ and p21^Waf1/Cip1^ and histone methylation of H3K4 in human CRC cells and suppress tumor growth *in vitro* and *in vivo* (Miyazaki et al., 2015; Ogawa et al., 2015).

In this study, we sought to clarify whether mir-200c, mir-302s and mir-369s would inhibit tumorigenesis in the colorectum of *CPC;Apc* mice, in which colon and rectal polyps are preferentially produced (Hinoi et al., 2007). We then performed microarray analysis to assess differential gene expression between normal colon mucosa and tumors from *CPC;Apc* mice. Based on the results, we focused on *c-MAF* (musculoaponeurotic fibrosarcoma) gene (Nishizawa et al., 1989).

The MAF family protein functions as a transcription factor of AP-1 family. c-MAF constitutes a large MAF family, together with MAFA and MAFB. It is reported that MAF may have tumor suppressive roles through regulating p53-dependent cell death, inhibition of MYB, induction of the cell cycle inhibitor p27^Kip1^ (Brundage et al., 2014). c-MAF is a human analogue of v-MAF which was identified as a retroviral oncoprotein in 1989, from avian retrovirus AS42 derived from a chicken musculoaponeurotic fibrosarcoma (Nishizawa *et al*., 1989).

To elucidate the role of c-MAF in CRC, we evaluated its expression in normal mucosa, early cancer, and advanced cancer samples from patients with CRC and performed i*n vitro* mechanistic studies in intestinal IEC-18 cells and CRC cell lines. Finally, we generated *c-MAF* knockout mice and investigated how tumor formation would be affected during chemical carcinogenesis with azoxymethane (AOM)/dextran sodium sulfate (DSS) treatment. Overall, our findings highlight c-MAF as a potential tumor suppressor in CRC.

## Results

### miRNA treatment suppresses colorectal tumor formation in *CPC;Apc* mice

We intravenously injected miR-200c-3p, miR-302-3p (-a,-b,-c,-d), and miR-369 (-3p, -5p) simultaneously (Miyoshi *et al*., 2011) (Miyazaki *et al*., 2015), or negative control (NC) miRNA (Supplementary Table S1) into the tail vein of *CPC;Apc* mice (Hinoi *et al*., 2007), three times weekly for 8 weeks, using the super carbonate apatite (sCA) delivery system (Fig. 1A; (Wu et al., 2015). Colorectal tumor burden was surveyed by rectosigmoid endoscopy at 9, 11, and 13 weeks (Fig. 1B) and directly confirmed postmortem at 15 weeks (Fig. 1C). The incidence of polyps in mice treated with the trio of miRNAs was significantly lower than in the NC animals (0.5 ± 0.2 vs. 3.3 ± 1.5 polyps/mouse, respectively; *P* = 0.026; Fig. 1D).

**Figure 1.**
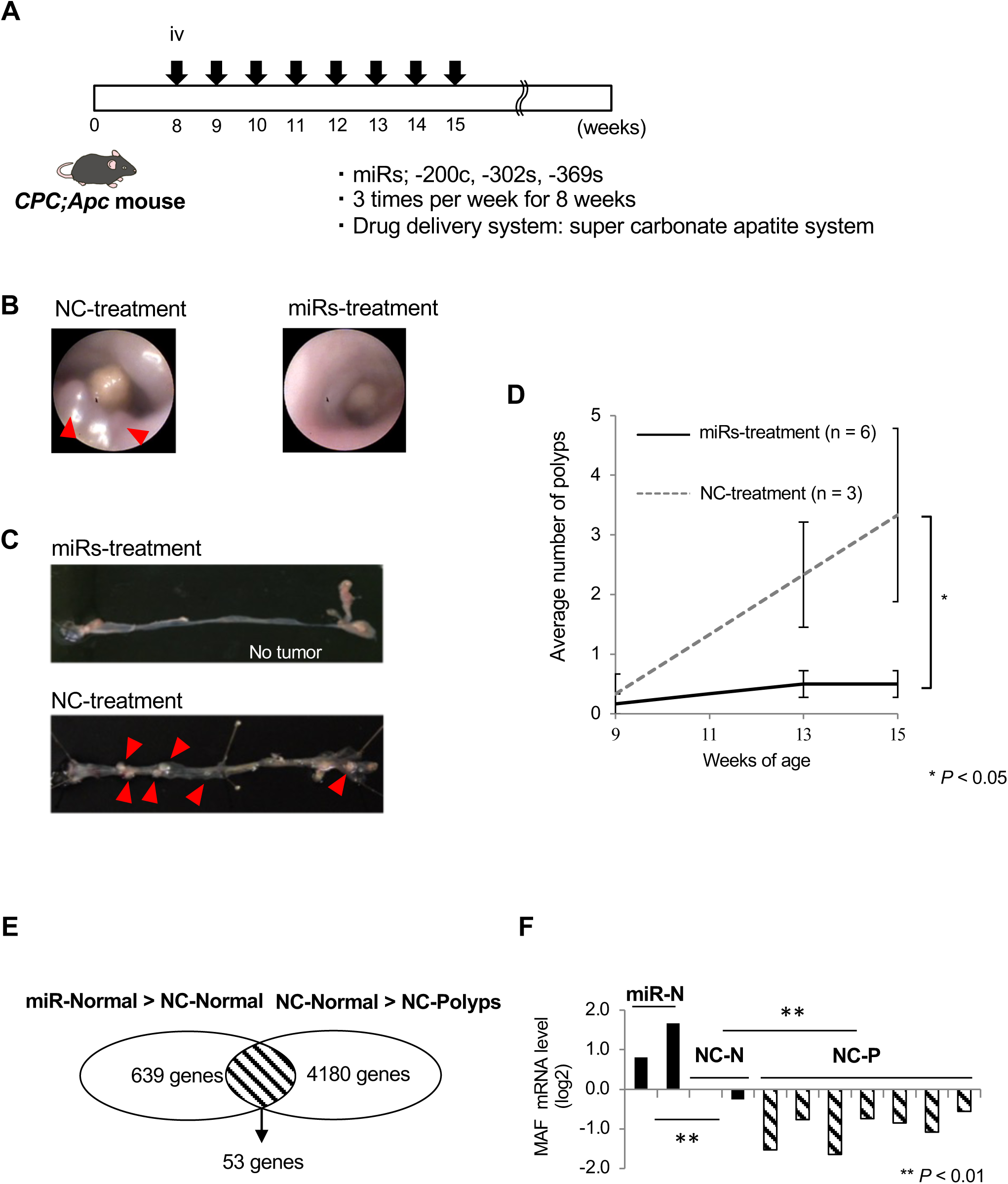
MicroRNA treatment suppressed colorectal tumor formation in *CPC;Apc* mice. (A) Experimental design. miR-200c-3p, miR-302-3p (-a,-b,-c,-d), and miR-369 (- 3p, -5p) or negative control (NC) miRNA was intravenously injected into the tail vein of *CPC;Apc* mice, three times per week for 8 weeks using the super carbonate apatite delivery system. (B) Images of distal colon observed by a rectosigmoid endoscopy at 13 weeks. (C) Mice were sacrificed at 15 weeks, and the colorectum was opened. Red arrowheads indicate polyp formation. (D) The incidence of polyps in mice treated with miRNAs was significantly lower than in negative control (NC)-treated mice (0.5 ± 0.2 vs. 3.3 ± 1.5 polyps/mouse, respectively; \**P* = 0.026). (E) Microarray analysis revealed that 53 genes in NC-treated normal mucosa were downregulated compared with the miRNA-treated normal mucosa and upregulated compared with NC-treated polyps. (F) qRT-PCR results indicated that c-MAF mRNA expression in the miRNA-treated normal mucosa was significantly higher than in control normal mucosa, and c-MAF mRNA was significantly decreased in polyps compared with control normal mucosa (\*\**P* < 0.001).

In a subset of samples, we next performed microarray analysis for differential mRNA expression between normal mucosa and colorectal polyps. Heat map analysis showed that 15 genes were highly expressed in polyps, and 53 genes showed stepwise downregulation from miRNA-treated normal mucosa to NC normal mucosa to NC polyps (Supplementary Fig. S1, Supplementary Table S2, Fig. 1E). Among these genes, we confirmed by qRT-PCR that c-MAF mRNA expression in the miRNA-treated normal mucosa was significantly higher than in NC normal mucosa, and that c-MAF mRNA was significantly decreased in polyps as compared with NC normal mucosa (*P* < 0.001 for both; Fig. 1F).

A database survey indicated that c-MAF mRNA expression was decreased in adenocarcinoma of the colon and rectum compared with normal mucosa (Supplementary Fig. S2A; (Rhodes et al., 2004). Moreover, several human malignancies, including colon and rectal cancer, have been found to express less c-MAF mRNA than normal tissues (Supplementary Fig. S2B [http://firebrowse.org/] and S2C).

### c-MAF expression in normal epithelial and CRC tissues

We found by qRT-PCR that c-MAF mRNA expression in CRC tissues was significantly lower than in paired normal mucosa samples (*P* = 0.045; Fig. 2A). Immunohistochemistry for c-MAF with duodenum samples as a positive control (Fig. 2B) showed intense nuclear staining of the c-MAF protein in normal mucosa from the bottom to the top of the glands. In contrast, we detected heterogenous c-MAF expression in advanced CRC tissue samples (Fig. 2C). High nuclear expression of the c-MAF protein in advanced CRC tissues was significantly decreased compared with expression in normal epithelium (*P* < 0.05; Fig. 2C), and this downregulation occurred at the early cancer stage (*P* = 0.038; Fig. 2D).

**Figure 2.**
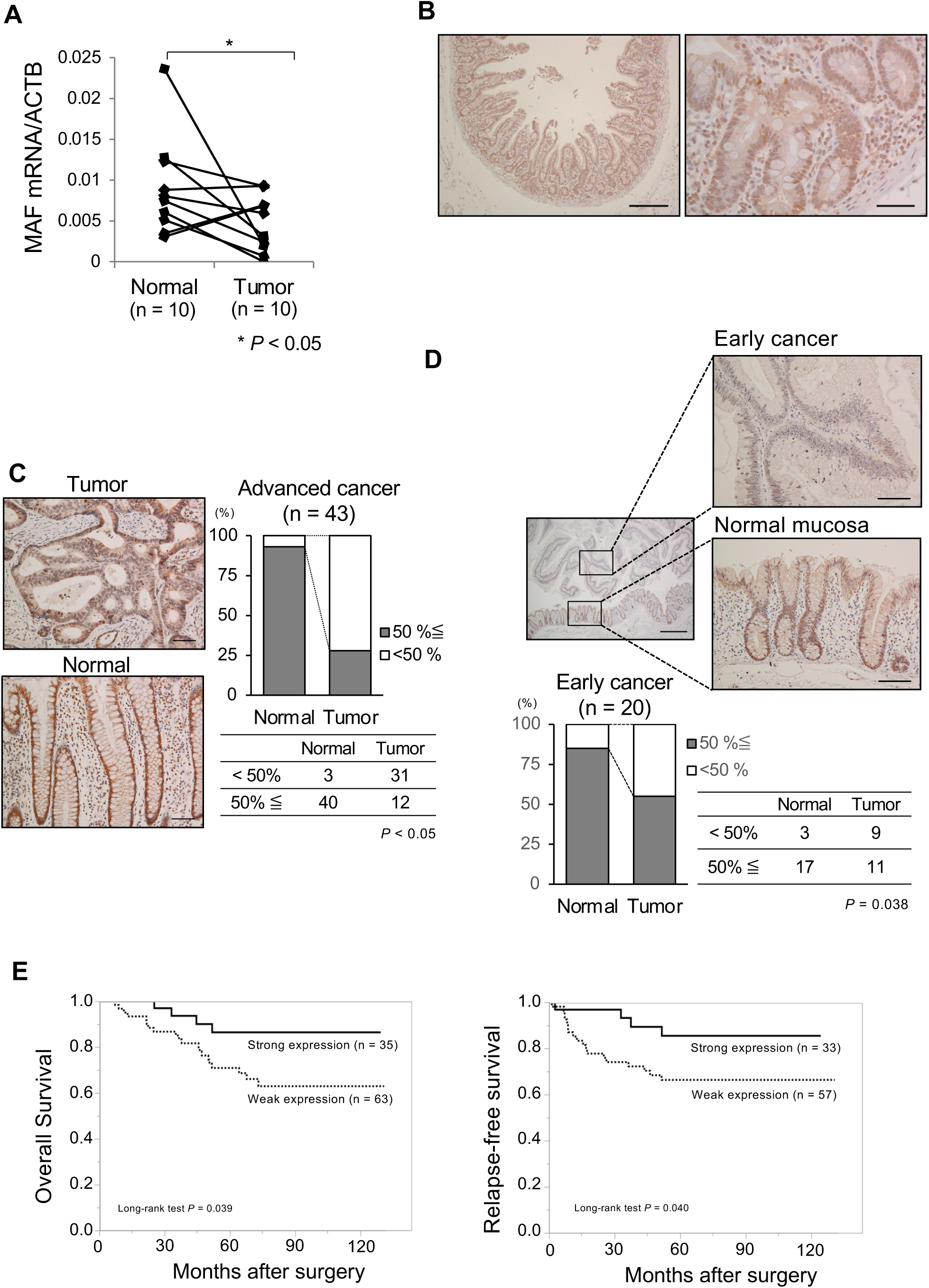
c-MAF expression in clinical samples of CRC. (A) c-MAF mRNA level in human CRC tissues was significantly lower than in their pair-matched adjacent normal mucosal tissues (*P* = 0.045). (B) Immunostaining of c-MAF protein in the duodenum as a positive control. Nuclear staining of c-MAF was noted in the duodenal epithelium. Scale bar: 100 μm. (C) Immunostaining of c-MAF protein in normal mucosa and advanced CRC tissue; depth of invasion defined as T2 (deep or deeper than the muscularis propria). c-MAF expression was noted in the nucleus of normal epithelium and tumor cells. When the c-MAF positive staining cutoff was set at >50%, the positive proportion in CRC tissues was significantly lower compared with normal mucosa (*P* < 0.05). Scale bar: 100 μm. (D) c-MAF expression in early cancer and adjacent normal mucosa; early cancer defined as T0 and T1 (invasion into lamina propria or submucosa). Cancer cells showed low c-MAF expression, whereas normal epithelial cells had strong c-MAF staining. Scale bars: left panel, 500 μm; right panels, 100 μm. Investigation of 20 early cancer samples revealed downregulation of c-MAF at an early cancer stage. (E) Setting c-MAF expression in normal mucosa; as a basis, we divided the CRC cases into two groups: strong (tumor > normal, n = 35) and weak (tumor < normal, n = 63). The Kaplan–Meier survival curve shows better prognosis for overall survival in the strong expression group (*P* = 0.039; median follow-up 66.4 [range 41.2– 791.8] months). Relapse-free survival was examined after exclusion of eight patients with stage IV disease, yielding a similar result (*P* = 0.040; median follow-up 63.5 [range 32.2–89.8] years).

Survival analysis in patients with advanced CRC revealed significantly prolonged overall survival (OS) in the group with strong c-MAF expression compared with those showing weak c-MAF expression (*P* = 0.039, median follow-up 66.4 months; Fig. 2E). Similarly, Kaplan–Meier survival curves showed significantly better relapse-free survival (RFS) in the group with strong c-MAF expression compared with the group showing low c-MAF expression (*P* = 0.040, median follow-up 63.5 months; Fig. 2E). A clinicopathological survey indicated that lymphatic duct invasion was significantly associated with weak c-MAF expression (*P* = 0.002; Supplementary Table S3). Multivariate analysis using a Cox proportional hazard model showed that c-MAF expression tend to be an independent prognostic factor for OS (hazard ratio [HR] 3.084, 95% confidence interval [CI] 0.921–10.320, *P* = 0.068) and that c-MAF was a significant independent indicator of better prognosis with RFS (HR 3.935, 95% CI 1.038–14.919, *P* = 0.044).

### Anti-tumor function of c-MAF in rat intestinal epithelial cells and CRC cells

To investigate the fundamental function of c-MAF, we performed knockdown experiments using siRNA. c-MAF mRNA expression significantly decreased after transduction of c-MAF siRNA into the IEC-18 rat intestinal cell line (Fig. 3A). Knockdown of c-MAF mRNA led to a significant increase in cell proliferation and colony-forming ability (*P <* 0.05 for both; Fig. 3B and C).

**Figure 3.**
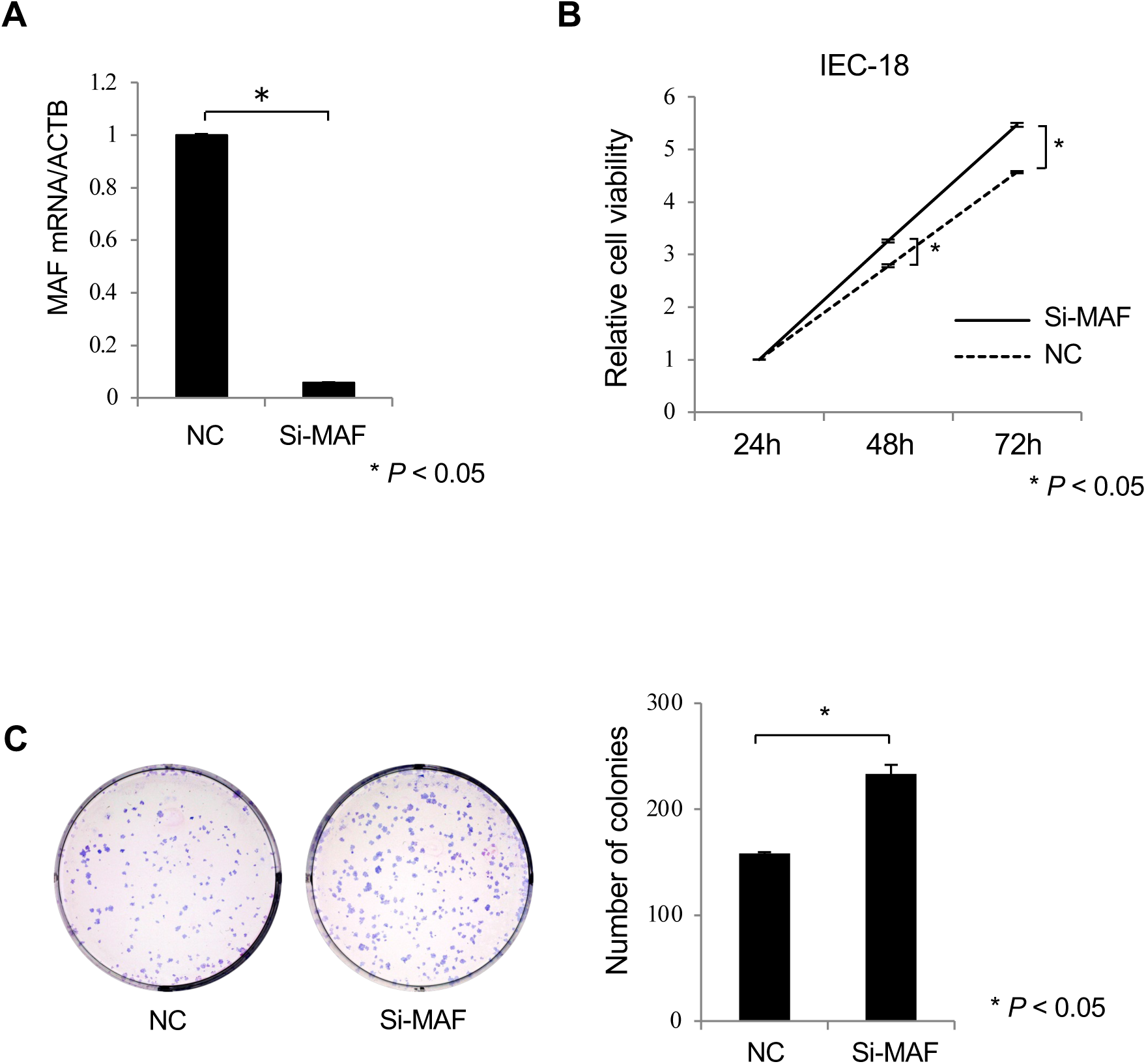
Small interfering (si)RNA knockdown of c-MAF in IEC-18 cells. (A) c-MAF siRNA or negative control siRNA was transfected into rat intestinal IEC-18 cells at 30 nM. c-MAF mRNA expression was measured by qRT-PCR. (B) Knockdown of c-MAF expression led to increased cell proliferation. (C) Colony-forming ability was significantly increased in c-MAF knockdown cells compared with negative control cultures. Left panel: representative pictures of Giemsa staining of colonies in the 6-well plate. Data are mean ± SD. Si-MAF, c-MAF-siRNA; NC, negative control siRNA. \**P* < 0.05.

We then transduced c-MAF cDNA into the HCT116 and LS174T CRC cell lines, both of which harbor wild-type *p53*. c-MAF–overexpressing CRC cells showed considerably higher c-MAF mRNA expression than empty vector (EV)-transduced control cells. In both cell lines, cell proliferation was significantly inhibited in the c-MAF–overexpressed cells compared with EV control cells (*P <* 0.05; Fig. 4A). In contrast, exogenous c-MAF transduction did not affect cell proliferation in *p53*-null HCT116 cells (Fig. 4B). In the HCT116 cells retaining wild-type *p53*, c-MAF overexpression induced cyclin-dependent kinase inhibitor p21^Waf1/Cip1^ expression as well as p53 expression at the RNA and protein levels, but did not do so in *p53*-null HCT116 cells (Fig. 4C and D). We also found that c-MAF overexpression enhanced the sensitivity to fluorouracil (5-FU) in HCT116 cells (Fig. 4E). Results of an Annexin V assay indicated that transduction of c-MAF significantly enhanced apoptosis with 24-h treatment with 100 μM 5-FU (*P* < 0.05; Fig. 4F).

**Figure 4.**
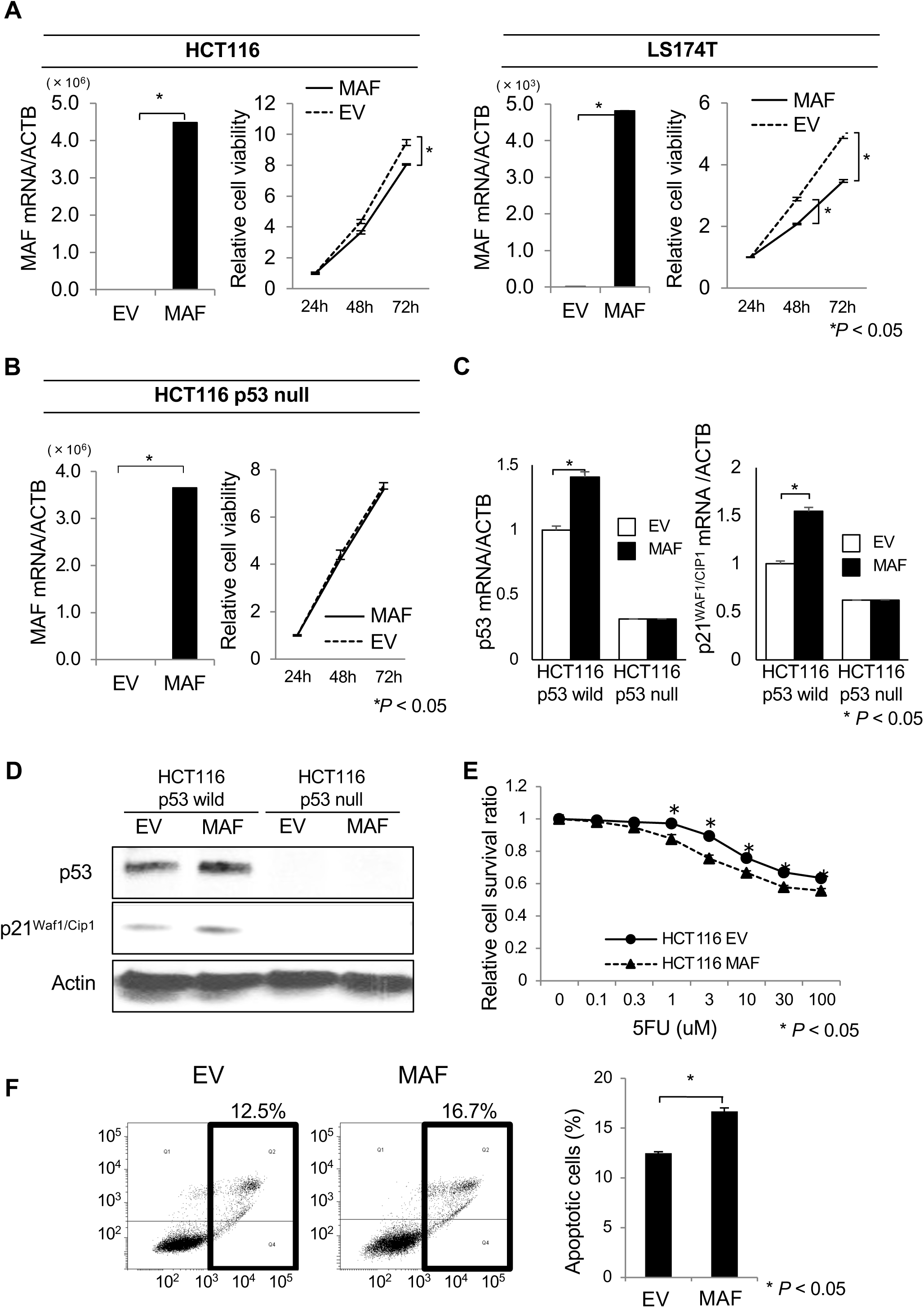
Effects of c-MAF overexpression on proliferative ability and 5-FU– induced apoptosis in CRC lines. (A) Overexpression of c-MAF in HCT116 and LS174T cells was validated by qRT-PCR. Cell proliferation of c-MAF–overexpressed cells (MAF) was decreased compared with empty vector (EV)–transduced control cells (n = 3 for each). (B) Overexpression of c-MAF in HCT116 p53-null cells was validated by qRT-PCR. Cell proliferation was not changed between c-MAF–overexpressed cells and control cells (n = 3). (C, D) Expression of p53 and p21^Waf1/Cip1^ in c-MAF– overexpressed HCT116 and HCT116 p53-null cells was measured by (C) qRT-PCR and (D) immunoblotting. Cells were collected 48 h after transfection. (E) Cells were harvested 24 h after transfection with either c-MAF expression vector (MAF) or empty vector control (EV) and seeded to 96-well plates for cytotoxicity assay under 5-FU treatment for 24 h. Sensitivity to 5-FU was significantly enhanced in c-MAF– overexpressed HCT116 cells (n = 6 for each concentration); \**P* < 0.05. (F) Annexin V apoptosis assay was performed with Annexin V and propidium iodide staining and flow cytometry analysis. c-MAF–overexpressed HCT116 cells showed increased 5-FU– induced apoptosis compared with vector control cells (n = 3 for each); \**P* < 0.05. Data are presented as mean ± SD.

### Effect of *c-MAF* knockout on colorectal tumor formation

To investigate the effect of c-MAF on carcinogenesis of the colorectum, we generated *c-MAF* KO mice in which one or eight nucleotides were deleted in 3’ flanking region of ATG transcription start codon (Supplementary Fig. S3A and B). Among 187 *c-MAF* KO mice and 37 wild-type mice, only 3 KO mice eventually developed tumors by 2 years after birth (Fig. 5A). A rectal adenocarcinoma arose in one *c-MAF* KO mouse bearing the eight-nucleotide depletion (hetero status) (Supplementary Fig. S4A). Tumor formation also was observed in the thorax and abdominal cavity in the other two mice. Based on immunohistochemistry results showing S100-positive and SOX10-negative cells, the tumor in the thorax was diagnosed as a malignant schwannoma (Supplementary Fig. S4B). The tumor in the abdominal cavity showed weakly positive staining for BCL6 and was positive for CD45R and was diagnosed as Burkitt’s lymphoma (Supplementary Fig. S4C).

**Figure 5.**
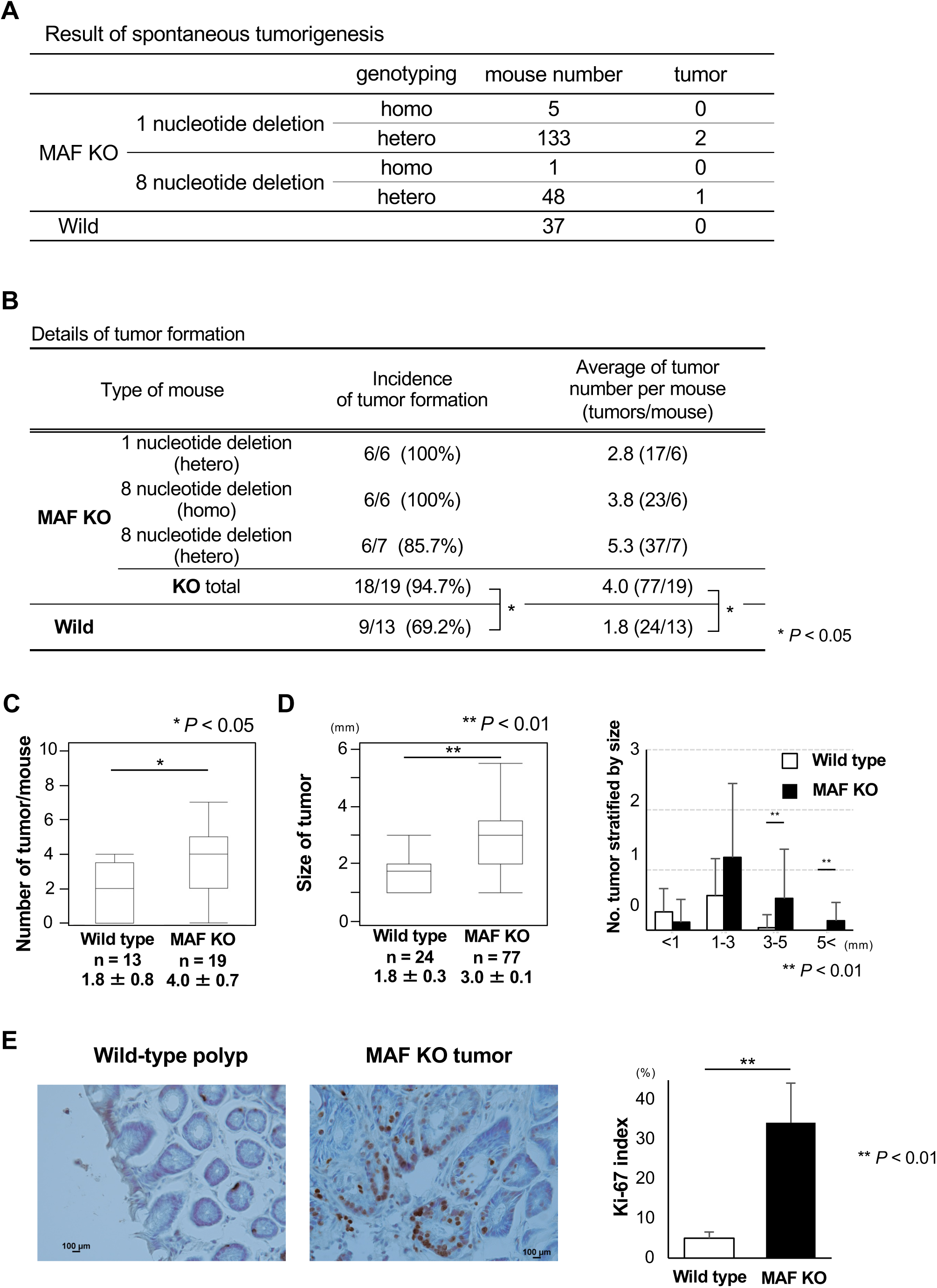
Spontaneous or colitis-associated tumor formation. (A) Tumors that spontaneously occurred are summarized. Only three tumors were generated in *c-MAF* KO mice around 2 years after birth. (B) Details of colitis-associated tumor formation; n = 32 mice (19 *c-MAF* KO and 13 wild type). The majority of *c-MAF* KO mice (18 of 19, 94.7%) developed colorectal tumors whereas 9 of 13 (69.2%) of wild-type mice did. The incidence of tumor formation was significantly higher in *c-MAF* KO mice; *\*P* < 0.05. The average tumor number in each group is also shown. (C) Number of tumors per mouse. *c-MAF* KO mice produced significantly more tumors than did wild-type mice (\**P* < 0.05). (D) Tumor size was significantly larger in *c-MAF* KO mice compared with wild-type mice (\*\**P* < 0.01). When stratified by tumor diameter at 1, 3, and 5 mm, c-MAF KO mice had significantly more tumor formation >3 mm (\*\**P* < 0.01; N.S., not significant). (E) Immunohistochemical staining for Ki-67 in tumors. *c-MAF* KO tumors had a significantly higher Ki-67 index as compared with wild-type tumors (\*\**P* < 0.01). Data are presented as mean ± SD.

### c-MAF deficiency increases tumor formation under chemical carcinogenesis

*c-MAF* KO mice (n = 19) and wild-type mice (n = 13) were treated with initial *i.p.* administration of 10 mg/kg AOM and two cycles of drinking water containing 2.0% DSS according to the protocol shown in Fig. 5B. As a whole, the tumors were positioned at the rectum and distal colon in either *c-MAF* KO or wild-type mice (Fig. 5C). In one *c-MAF* KO mouse, a rectal tumor prolapsed from the anus approximately 100 days after *i.p.* injection of AOM (Fig. 5D, I–III (a)). Hematoxylin and eosin (H&E) staining revealed a moderately differentiated adenocarcinoma (Fig. 5D, IV-V).

Figure 5E summarizes tumor formation, which was significantly higher in *c-MAF* KO than in wild-type mice (*P* < 0.05). *c-MAF* KO mice produced significantly more tumors than did wild-type mice (1.8 ± 0.8 vs. 4.0 ± 0.7, *P* < 0.05; Fig. 5E and F). Regarding tumor size, *c-MAF* KO mice had significantly larger tumors than their wild-type counterparts (*P* < 0.01; Fig. 5G), as was especially evident for tumors larger than 3 mm (*P* < 0.01; Fig. 5G). The Ki-67 index, a proliferation marker, was significantly higher in *c-MAF* KO than wild-type tumors (*P* < 0.01; Fig. 5H).

### Comparative gene expression analysis of AOM/DSS-treated *c-MAF* KO and wild-type mice

To elucidate the underlying mechanism by which AOM/DSS treatment resulted in a significant increase in tumor formation in *c-MAF* KO mice, we performed RNA sequencing (wild-type normal, n = 4; wild-type tumor, n = 9; *c-MAF* KO normal, n = 6; *c-MAF* KO tumor, n = 5). The heat map showed numbers of downregulated or upregulated gene expression between *c-MAF* KO and wild-type mice in normal mucosa or tumors (Supplementary Fig. S5A and B). Ingenuity Pathway Analysis of normal mucosa indicated that many cancer-promoting growth factors, transcription factors, kinases, and cytokines were activated as the upstream regulators in *c-MAF* KO mice (Table 2), as also was the case with tumor samples from *c-MAF* KO mice (Supplementary Table S4). Downstream analysis for disease and function showed that many cancer-related factors were activated in *c-MAF* KO tumors (Supplementary Fig. S6).

**Table 1.** Univariate and multivariate analysis of clinicopathological characteristics associated with overall survival and relapse-free survival.

| Univariate and multivariate analysis of clinicopathological characteristics associated with OS<sup>a</sup> |  |  |  |  |  |  |
| --- | --- | --- | --- | --- | --- | --- |
|  | Univariate |  |  | Multivariate |  |  |
|  | HR<sup>b</sup> | 95% CI<sup>c</sup> | *P* value | HR | 95% CI | *P* value |
| Age ($\geq 65$/$< 65$) | 3.331 | 1.319-8.411 | 0.011 | 4.37 | 1.461-13.072 | 0.008 |
| Gender (female/male) | 0.642 | 0.275-1.501 | 0.307 |  |  |  |
| Location (rectum/colon) | 0.405 | 0.161-1.020 | 0.055 | 0.250 | 0.082-0.764 | 0.015 |
| Depth (T4/Tis, T1, T2, T3) | 2.34 | 0.799-6.851 | 0.121 | 1.875 | 0.493-7.122 | 0.356 |
| Lymph node metastasis<br>(positive/negative) | 2.649 | 1.186-5.917 | 0.018 | 1.544 | 0.608-3.923 | 0.361 |
| Histological type<br>(por, sig, muc/tub1, tub2) | 3.204 | 1.093-9.396 | 0.034 | 7.830 | 2.000-30.651 | 0.003 |
| Lymphatic duct invasion<br>(positive/negative) | 3.173 | 0.946-10.644 | 0.062 | 1.398 | 0.334-5.857 | 0.646 |
| Venous invasion<br>(positive/negative) | 2.666 | 1.195-5.944 | 0.017 | 1.971 | 0.763-5.094 | 0.161 |
| MAF expression<br>(weak/strong) | 2.934 | 1.002-8.585 | 0.049 | 3.084 | 0.921-10.320 | 0.068 |
| Univariate and multivariate analysis of clinicopathological characteristics associated with RFS<sup>d</sup> |  |  |  |  |  |  |
|  | Univariate |  |  | Multivariate |  |  |
|  | HR | 95% CI | *P* value | HR | 95% CI | *P* value |
| Age ($\geq 65$/$< 65$) | 6.355 | 1.878-21.502 | 0.003 | 10.464 | 2.693-40.659 | <<br>0.001 |
| Gender (female/male) | 0.795 | 0.333-1.895 | 0.604 |  |  |  |
| Location (rectum/colon) | 0.510 | 0.208-1.251 | 0.141 | 0.432 | 0.165-1.133 | 0.088 |
| Depth (T4/Tis, T1, T2, T3)<sup>e</sup> | 4.081 | 1.502-11.092 | 0.006 | 6.388 | 1.570-25.989 | 0.010 |
| Lymph node metastasis<br>(positive/negative) | 2.787 | 1.207-6.436 | 0.016 | 1.853 | 0.648-5.299 | 0.250 |
| Histological type (por, sig,<br>muc/tub1, tub2)<sup>f</sup> | 1.441 | 0.336-6.177 | 0.623 |  |  |  |
| Lymphatic duct invasion<br>(positive/negative) | 2.544 | 0.861-7.520 | 0.091 | 1.124 | 0.286-4.416 | 0.867 |
| Venous invasion<br>(positive/negative) | 2.697 | 1.163-6.253 | 0.021 | 1.341 | 0.444-4.051 | 0.602 |
| MAF expression<br>(weak/strong) | 2.945 | 0.996-8.706 | 0.051 | 3.935 | 1.038-14.919 | 0.044 |

a) OS, overall survival; b) HR, hazard ratio; c) CI, confidence interval; d) RFS, relapse-free survival; e) Tis, carcinoma in situ; T1, involvement of submucosa; T2, involvement of muscularis propria; T3, involvement of subserosa; T4, involvement of serosal surface or direct invasion to other organs; f) tub1, well differentiated adenocarcinoma; tub2, moderately differentiated adenocarcinoma; muc, mucinous carcinoma; por, poorly differentiated adenocarcinoma

**Table 2.**
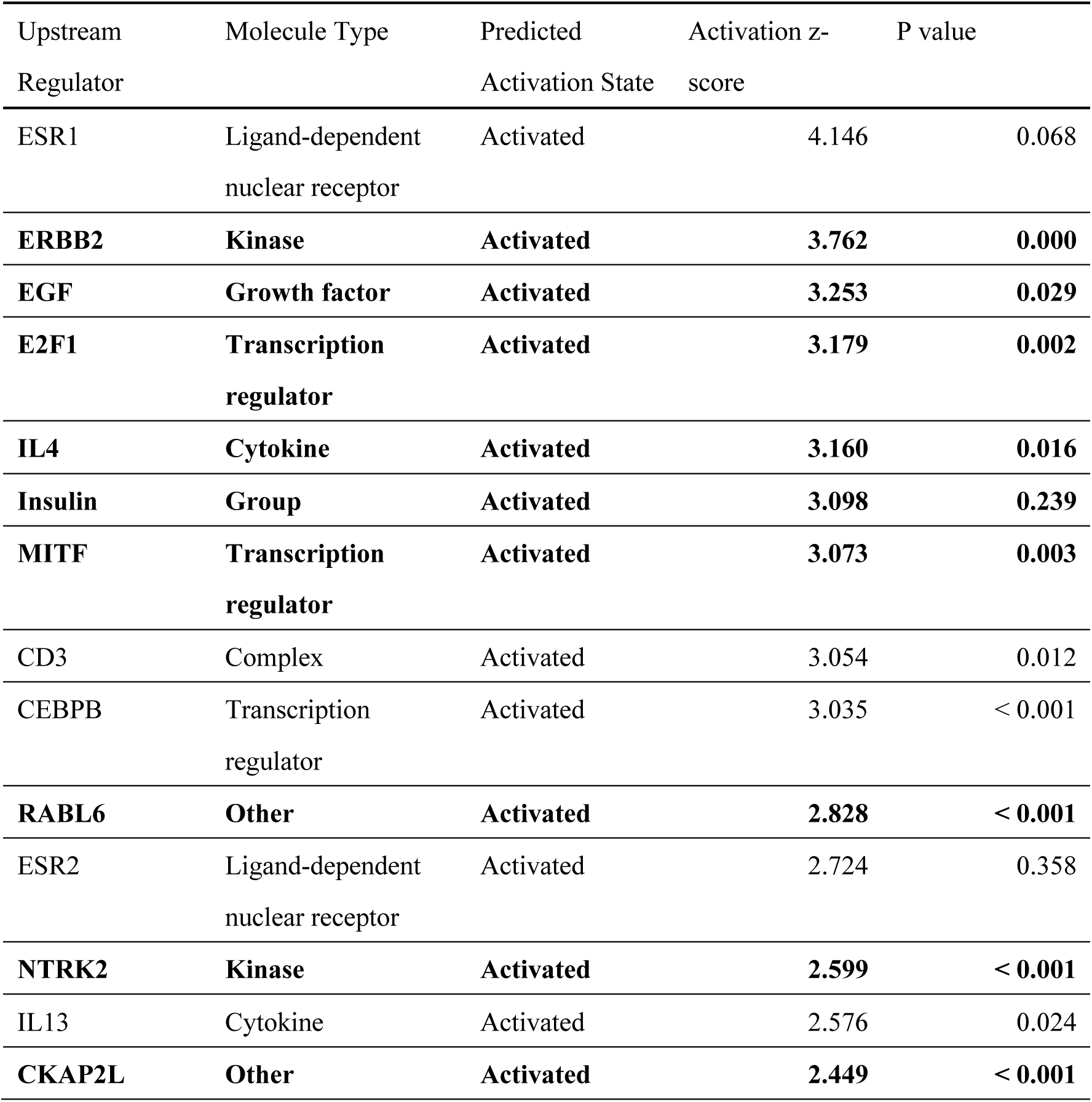
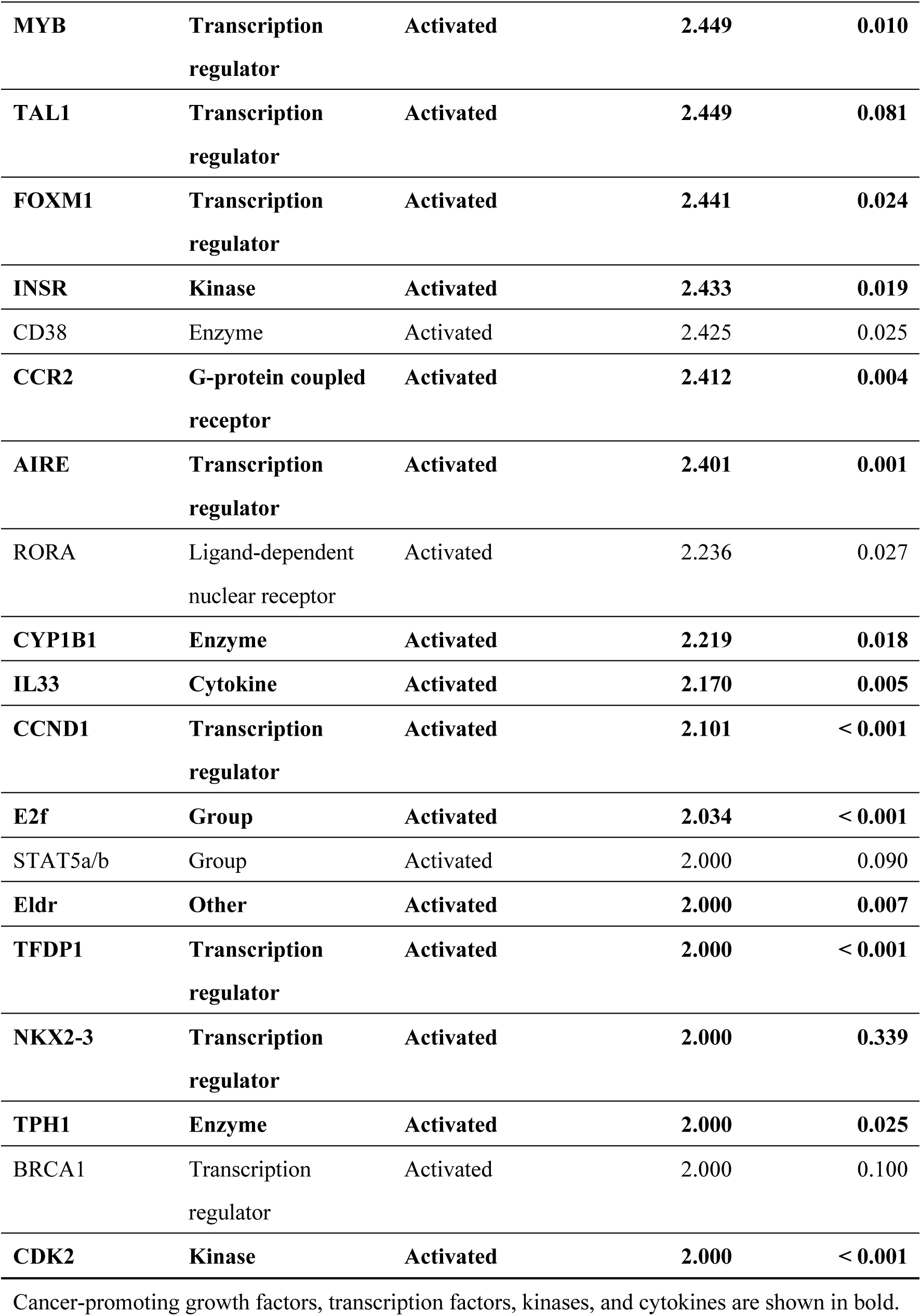
Lists of activated molecules in normal mucosa of *c-MAF* KO mice compared with wild-type mice after treatment with AOM/DSS. Ingenuity Pathway Analysis of normal mucosa showed that many cancer-promoting growth factors, transcription factors, kinases, and cytokines were activated as upstream regulators in *c-MAF* KO mice.

### Inverse relationship between c-MAF protein and p53 protein expression

We then performed comparative immunohistochemical analysis for c-MAF and p53 protein because the c-MAF transcription factor can activate *p53* transcription (Hale et al., 2000). We found an inverse staining pattern between c-MAF and p53 expression (Fig. 6A, Case A and Case B). Of note, this reciprocal staining pattern was found even within identical CRC tissue samples (Fig. 6B), so that cells that accumulated p53 protein lost c-MAF expression, and vice versa. As a whole, we found an inverse relationship between c-MAF and p53 expression in CRC tissue samples when the cutoff point was set at 10% for p53 positivity (*P* = 0.024; Fig. 6C). When we transduced two types of mutant *p53* (R175H and R248W) into p53-null HCT116 cells, the clones displayed decreased c-MAF expression and increased miR-155 (Supplementary Fig. S7A-C). Previous studies had shown that miR-155 directly bound to the 3’-untranslated region of c-MAF (Rodriguez et al., 2007) (Su et al., 2014) (Wolf et al., 2013), and we confirmed that transfection of mature miR-155 suppressed c-MAF expression (Supplementary Fig. S7D).

**Figure 6.**
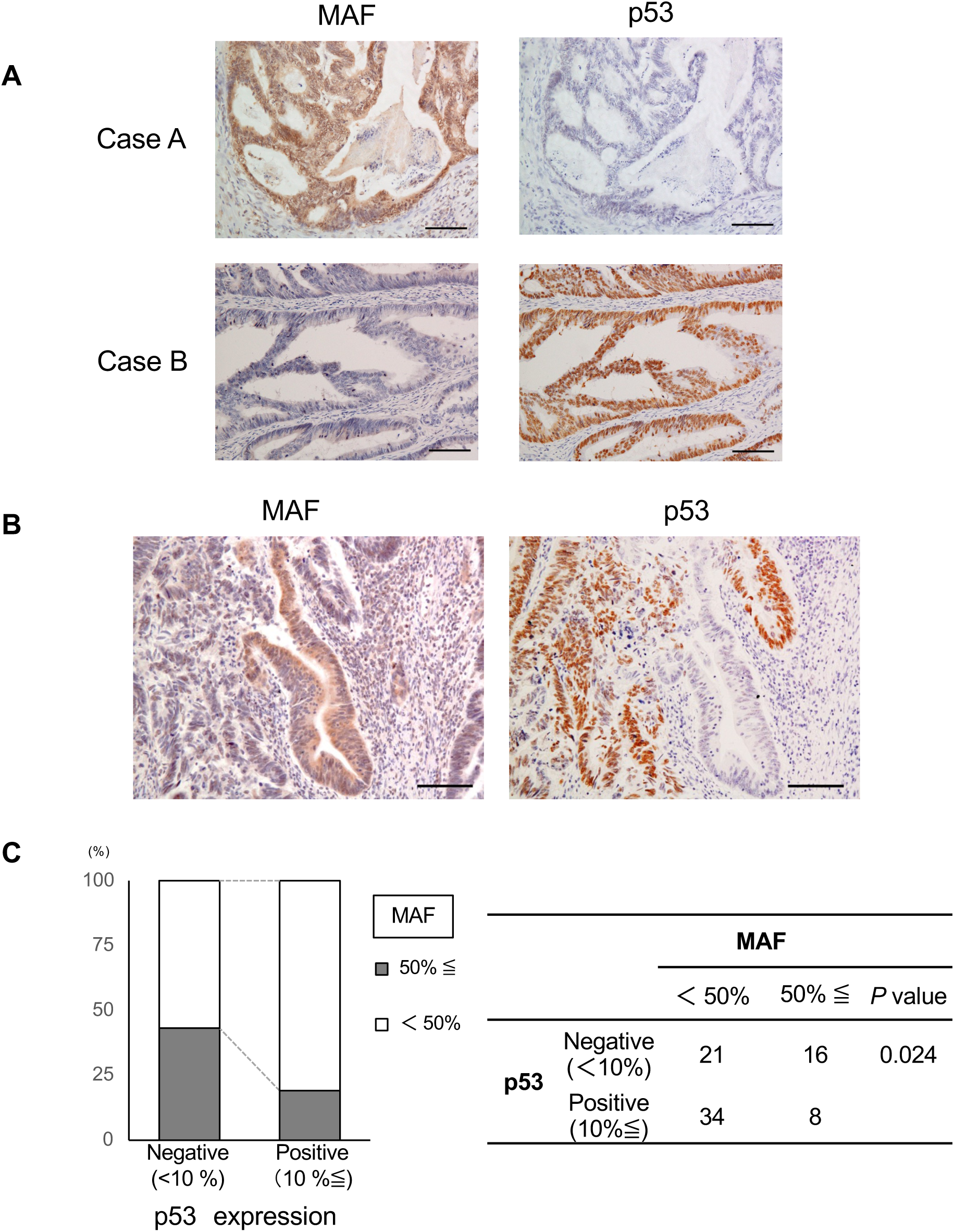
Inverse relationship between c-MAF and p53 protein expression. (A) Representative pictures of p53 staining in two human CRC tissue samples. Left: c- MAF; right, p53. c-MAF and p53 showed complementarily positive findings. (B) A set of pictures showing an inverse staining pattern between c-MAF and p53 expression in serial sections. (C) A significantly inverse correlation was observed between the expression of c-MAF and p53 (*P* = 0.024).

Because c-MAF expression might be indirectly upregulated through a complex reaction among the miRNAs, we focused on one of several possible candidate pathways, the GATA3-cMAF axis (Naito et al., 2011). To assess this route, we examined differential gene expression in human MRC5 lung fibroblasts transfected with human miR-200c, miR-302s, and miR-369s simultaneously. RNA sequencing indicated upregulation of GATA3 mRNA by 3.07-fold in transfected compared with control cells (*P* = 0.035; Supplementary Fig. S8A, data deposited at https://www.ncbi.nlm.nih.gov/geo/query/acc.cgi?acc=GSE210980). Ingenuity Pathway Analysis identified GATA3 as an upstream activator of c-MAF (Supplementary Fig. S8A). When we introduced human miR-200c, miR-302s, and miR- 369s simultaneously into MRC5 lung fibroblasts and HBEC3-KT bronchial epithelial cells, qRT-PCR showed increased expression of GATA3 and c-MAF mRNA (Supplementary Fig. S8B).

## Discussion

miRNAs are involved not only in progression of human cancers but also in carcinogenesis. It is reported that miR-26a can overcome potential oncogenic activity in intestinal tumorigenesis of *Apc^min/+^*mice and that tumor formation is abundant in miR- 10a–deficient mice (Stadthagen et al., 2013; Zeitels et al., 2014). We previously reported an inhibitory effect of administering mixture of miR-200c, miR-302s, and miR-369s on *in vivo* tumor growth of CRC cells (Miyazaki et al., 2015; Ogawa et al., 2015). In this study, we employed *CPC;Apc* mice because polyp formation can be readily monitored using periodical rectosigmoid endoscopy, and found that systemic administration of the miRNAs simultaneously suppressed tumor formation in the colorectum of these animals. Of note, our approach using intravenous injection of the miRNAs steps around virus-derived genome integration, so that it is safe and suitable for clinical application, and we and others have confirmed its efficacy in animal studies (Abd-Aziz et al., 2020; Forterre et al., 2020; Fukata et al., 2018; Hiraki et al., 2015; Inoue et al., 2018; Merhautova et al., 2016; Morimoto et al., 2020; Takahashi et al., 2019; Takeyama et al., 2014; Tamai et al., 2018; Wang et al., 2022; Wu et al., 2015).

To explore the underlying mechanism of how the miRNAs suppress tumor formation, we analyzed expression of genes that were upregulated in miRNA-treated normal mucosa and downregulated in control tumors compared with NC normal mucosa. Of 53 candidate genes, c-MAF attracted our interest because its bird homologue v-MAF is considered an oncogene (Nishizawa et al., 1989) and its increased mRNA expression and an association with malignant properties have been reported in multiple myeloma (Hurt et al., 2004) and T-cell lymphoma (Morito et al., 2006), which is paradoxical to our initial results. In addition, MAF has been reported to be a mediator of breast cancer bone metastasis through regulating parathyroid hormone–related protein (Pavlovic et al., 2015). According to a public database, however, human c-MAF mRNA expression is downregulated in carcinomas of the cervix and uterus, colorectum, stomach, bile duct, breast, bladder, and organs (http://firebrowse.org/). Based on these findings, we hypothesized that *c-MAF* could function as either an oncogene or a tumor suppressor, depending on cancer type, and made it the focus of our further investigation.

How the three miRNAs led to induction of c-MAF mRNA is unclear. Searches of public databases (TargetScan human version 7.2, miRwalk, miRabel, miRmap) did not turn up direct binding sites between miR-200c, miR-302s, and miR-369s and the 3’ untranslated region of c-MAF mRNA. Given the potential for c-MAF expression to be upregulated indirectly because of interactions among the miRNAs, we chose a candidate pathway, the GATA3–cMAF axis, to examine (Naito *et al*., 2011). Sequencing of RNA from MRC5 lung fibroblasts transfected with human miR-200c, miR-302s, and miR-369s simultaneously showed a more than 3-fold upregulation of GATA3 mRNA in transfected cells compared with control cells. Furthermore, GATA3 and c-MAF mRNA expression were increased in both MRC5 lung fibroblasts and HBEC3-KT bronchial epithelial cells after transfection with the three miRNAs. Pathway analysis identified GATA3 as an upstream activator of c-MAF.

Although c-MAF has been investigated in some cancers (Eychène et al., 2008; Hurt *et al*., 2004; Morito *et al*., 2006; Pavlovic *et al*., 2015), the expression profile and function of c-MAF in human cancers are still largely unknown. Moreover, our database survey showed that c-MAF mRNA expression was downregulated more often than not in a series of human malignancies including CRC, as shown in Supplementary Fig. S2A-C. At the protein level, we confirmed by immunohistochemistry an intense expression of c-MAF in colonic epithelial cells, with a decrease in samples from early stage CRC and a poor prognosis associated with low expression. Overall, our findings in *CPC;Apc* mice and clinical CRC samples suggest that c-MAF as a transcription factor may have a tumor-suppressive role in the generation and progression of CRC.

Our *in vitro* mechanistic studies showed that silencing c-MAF expression resulted in increased cell proliferation and enhanced colony-forming ability in the IEC- 18 intestinal cell line, further highlighting a tumor-suppressive role of c-MAF in intestinal cells. c-MAF binds the MAF recognition element in the mouse *p53* promoter and causes cell death through induction of the p53 protein. This region is conserved between mouse and human (Hale et al., 2000). *p53* is a predominant gene in controlling apoptosis in response to abnormal cell proliferation and stress (Benchimol, 2001; Levine, 1997) and may cooperate with c-MAF to exert a tumor-inhibitory effect. Using CRC cell lines, we showed that overexpression of c-MAF led to reduced cell proliferation and enhancement of 5-FU–dependent apoptosis in CRC cell lines harboring wild-type *p53*. Overexpression of c-MAF indeed increased the expression of p53 and its downstream target p21^waf1/cip1^, a negative cell cycle regulator (Abbas and Dutta, 2009; el-Deiry et al., 1994; el-Deiry et al., 1993) in the HCT116 cells with wild-type *p5*3. In contrast, forced expression of c-MAF did not induce p53 or p21^waf1/cip1^ and had no effect on cell proliferation in p53-null HCT116 cells. These findings suggest that c-MAF may play an anti-oncogenic role through p53 upregulation in intestinal and CRC cells.

In our *in vivo* experiment using *c-MAF* KO mice, we rarely observed spontaneous tumor formation, with an incidence of 3/187 (1.6%), including one rectal adenocarcinoma. Therefore, c-MAF alone is unlikely to be a definitive tumor-suppressor gene. However, our chemical carcinogenesis experiment made it clear that c-MAF supports suppression of tumor formation: *c-MAF* KO mice had a significantly higher incidence of colorectal tumors and larger tumor size, with a higher Ki-67 index. Ingenuity Pathway Analysis further demonstrated that *c-MAF* KO normal mucosa bears higher potential for tumor formation via activation of genes that reportedly promote carcinogenesis (Table 2A). For example, ERBB2 is an oncogene amplified in breast cancer (Calogero et al., 2007; Ursini-Siegel et al., 2007); E2F1 facilitates carcinogenesis in liver, brain, skin, and testis (Agger et al., 2005; Conner et al., 2000; Olson et al., 2007; Pierce et al., 1998); and TFDP1, a heterodimer partner of E2F, is reported to facilitate carcinogenesis in skin tissue (Wang et al., 2001). Other molecules are reported to have a role in carcinogenesis, such as MITF in kidney angiomyolipoma, Myb and FOXM1 in colon cancer, TAL-1 in T-cell acute lymphoblastic leukemia, and NKX2-3 in B-cell lymphoma (Condorelli et al., 1996; Malaterre et al., 2016; Robles et al., 2016; Yoshida et al., 2007; Zarei et al., 2021). A comparison of *c-MAF* KO tumors and wild-type tumors showed that *c-MAF* KO tumors exhibited activation of many tumor-promoting growth factors, such as VEGF, IGF1, HGF, EGF, TGFβ1, and FGF2, as well as the transcription factors Jun and STAT3, which activate signal transduction for tumor growth and survival (Fang and Richardson, 2005; Yu et al., 2014). Taken together, our data suggest that c-MAF behaves like a tumor suppressor in tumorigenesis of CRC.

c-MAF also may coordinate cells in differentiating into retina, sensory nerve, and immune T cells and maintain cell quiescence in the lens. This transcription factor thus also is considered to induce gene expression during the tissue-specific differentiation process (Andris et al., 2017; Eychène et al., 2008; Kataoka et al., 1994; Wende et al., 2012). Brundage *et al*. showed that c-MAF expression was downregulated in a malignant peripheral nerve sheath tumor cell line and that c-MAF suppressed cell proliferation and anchorage-independent growth, and induced cell differentiation and apoptosis (Brundage et al., 2014). They also observed that c-MAF paradoxically promoted *in vivo* tumor growth of NF1 (neurofibromatosis type 1) patient-derived malignant peripheral nerve sheath tumor cells (Brundage et al., 2014). These findings suggest that as a transcription factor, c-MAF may cooperate with optimal downstream targets according to the cell context, surrounding microenvironment, and/or cell type, so that it can act either as an oncogene or anti-oncogene.

The mechanism of how *c-MAF* is downregulated in CRC remains to be addressed. A few studies have described mechanisms of MAF regulation through deletion, loss-of-function mutations, and promoter methylation (Ivascu et al., 2007; Johnson and Fleet, 2013; Perveen et al., 2007), but such modifications are not reported in CRC. Here, we observed impressive staining results showing a reciprocal expression pattern between c-MAF and p53 in CRC tissue samples (Fig. 6B). This pattern implied that the role of c-MAF could extend beyond mouse or cell culture systems and be relevant in clinical CRC tissues. By immunohistochemistry, we found that the wild-type p53 protein was basically undetectable; therefore, a c-MAF–positive/p53-negative CRC pattern seems to make sense considering that c-MAF is a transcription factor that positively regulates *p53* (Fig. 6A, Case A). On the other hand, a mutated *p53* product is known to be detectable in the nucleus because of its prolonged half-life (Finlay et al., 1988). Of considerable interest is that c-MAF somehow lost its expression when mutated p53 protein was accumulated in the CRC cells (Fig. 6A, Case B; Fig. 6B). One possible underlying mechanism could be related to the mutant p53–miR-155-c–MAF axis.

Neilsen et al. reported that transduction of mutant *p53* upregulated miR-155 expression through p63 in breast cancer (Neilsen et al., 2013). miR-155 is one of the representative oncomirs and targets c-MAF by direct binding (Rodriguez et al., 2007; Su et al., 2014; Wolf et al., 2013). We confirmed this scenario by transduction of mutated *p53* into p53- null HCT116 CRC cells (Supplementary Fig. S7). *p53* is an important tumor suppressor acting at a critical transition point from adenoma to cancer (Ohue et al., 1994). It is assumed that p53 mutation could be one reason for c-MAF inhibition and that mutated p53-mediated abolition of c-MAF may further accelerate carcinogenesis and progression of CRC. However, we should emphasize that c-MAF could have a tumor-suppressive effect via mechanisms other than p53; some studies have shown that *p53* gene mutation is not detectable in AOM/DSS-induced CRC (De Robertis et al., 2011; Tanaka et al., 2003), whereas *c-MAF* KO revealed many other candidate factors facilitating carcinogenesis, as we show in Table 2A.

Taken together, the present findings imply a tumor-suppressive role of c-MAF in tumorigenesis and progression of CRC. Our data would provide c-MAF as a novel marker for prognosis of CRC patients and may lead to development of novel therapeutic option against CRC.

## Acknowledgements

HCT116 and p53-null derivatives were kind gifts from Prof. Bert Vogelstein (Johns Hopkins University, Baltimore, MD, USA). We are grateful to other collaborators Y. Kotani for generating c-MAF knockout mice, and H. Ogawa, M. Konno, M. Nomura, A Eto, K Kitagawa, A. Toyama, and K. Asai for their support in animal experiments, and genotyping.

This work was supported by JSPS Grant-in-Aid for Exploratory Research Grant Number 26670605 to H. Yamamoto, JSPS KAKENHI Grant Number 20K08356 to D. Okuzaki and Research Grants of Princess Takamatsu Cancer Research Funds 2017 to H. Yamamoto.

## Author contributions

HI and HY designed the study. T.Hata, YD, HE, and MM supervised the study. HY, YY, TA and T.Hinoi are responsible for methodology. HI, T.Hata, YY and HY analyzed and interpreted the data and confirm the authenticity of all the raw data. HI, T.Hata, DO, KT and KI performed the experiments. HI, YY and HY are responsible for the statistical analysis. TO, NM, HT, MU and TM collected and provided the normal and colorectal cancer tissue samples and their clinical data. HI wrote the original draft. HY, HT and YY reviewed and edited the manuscript. HI, KT, YM, QY, T.Hata and HH performed animal experiments. All authors have read and approved the final manuscript.

## Declaration of interests

The authors have no conflict of interest to disclose.

## Material and Methods

### Cell Lines

Human lung fibroblast (MRC5), human bronchial epithelial (HBEC3-KT), and human CRC (LS174T) cell lines and a rat intestinal cell line, IEC-18, were obtained from the American Type Culture Collection (Rockville, MD, USA). HCT116 p53^+/+^ cells retained the wild-type *p53* gene, whereas both alleles of the *p53* gene were deleted through homologous recombination in HCT116 p53^−/−^ cells. This genetically impaired HCT116 cell line and the parental line with wild-type genes were generous gifts from Dr. Bert Vogelstein (Johns Hopkins University School of Medicine, Baltimore, MD, USA). HBEC3-KT cells were maintained in Airway Epithelial Cell Basal Medium (ATCC PCS-300-030) supplemented with Bronchial Epithelial Cell Growth Lit (ATCC PCS-300-040). Other cell lines were maintained in Dulbecco’s modified Eagle medium (Sigma-Aldrich, St. Louis, MO, USA) containing 10% fetal bovine serum, 100 U/mL penicillin, and 100 μg/mL streptomycin. Cultures were maintained at 37°C in a humid incubator with 5% CO2.

### Chemicals

5-FU was purchased from Nacalai Tesque Inc. (Kyoto, Japan). AOM saline was purchased from Sigma-Aldrich. DSS was purchased from MP Biomedicals (Santa Ana, CA, USA).

### siRNA and miRNA

siRNA for rat-c-MAF (4390771) and its negative control siRNA (4390843) were purchased from Thermo Fisher Scientific (Waltham, MA, USA). miR- 155 mimic and its negative control sequence, and mouse and human miR-200c-3p, 302a-3p, 302b-3p, 302c-3p, 302d-3p, 369-3p, and 369-5p mimics and their negative control sequences were purchased from Gene Design, Inc. (Osaka, Japan). The sequence information is shown in Supplementary Table S1. Lipofectamine RNAiMax (Thermo Fisher Scientific) was used for transfection of siRNA or miRNA according to the manufacturer’s instructions.

### Plasmid DNA

pCMV6-c-MAF plasmid DNA was purchased from OriGene (Rockville, MD, USA). pCMV6-empty vector was used as a control. pCMV-Neo-Bam p53 R175H (R175H), pCMV-Neo-Bam p53 R248W (R248W), and pCMV-Neo-Bam (Empty) were purchased from Addgene (a gift from Bert Vogelstein, addgene, #16436; http://n2t.net/addgene:16436; RRID:Addgene_16436, #16437; http://n2t.net/addgene:16437; RRID:Addgene_16437, #16440; http://n2t.net/addgene:16440; RRID:Addgene_16440, respectively)(Baker et al., 1990). Transfection was performed with Lipofectamine 3000 Reagent (Thermo Fisher Scientific) according to the manufacturer’s protocol.

### Western Blot Analysis

Western blotting was performed as described previously (Hamabe et al., 2014). Briefly, cells were collected and lysed in RIPA buffer containing phosphatase inhibitor and protease inhibitor cocktail. The protein lysates (20 μg) from each sample were separated with sodium dodecyl sulfate-polyacrylamide gel electrophoresis and transferred onto a polyvinylidene difluoride membrane. The membrane was blocked with 5% skim milk and incubated with the primary antibody at a concentration of 1-2 μg/mL, as follows: anti-human c-MAF polyclonal antibody (ab72584, Abcam, Cambridge, UK), anti-human p53 monoclonal antibody (M7001, Dako, Santa Barbara, CA, USA), anti-human p21^Waf1/Cip1^ monoclonal antibody (ab80633, Abcam), and anti-human ACTB polyclonal antibody (A2066, Sigma-Aldrich). The membrane was incubated with secondary antibodies and visualized with the ECL Detection System (GE Healthcare, Little Chalfont, UK).

### Proliferation Assay

Cells were seeded at a density of 3–5 × 10^3^ cells per well in 96- well plates. After culture for 24, 48, or 72 h, cell viability was determined using the Cell Counting Kit-8 (Dojindo, Kumamoto, Japan) by measuring the absorbance at 450 nm using an iMarkTM microplate absorbance reader (Bio-Rad, Hercules, CA, USA).

### Colony-formation Assay

Cells were seeded at a density of 500 cells per well in a 6- well plate. After incubation at 37°C for 10 days, cells were washed with phosphate-buffered saline, fixed with 10% formalin, and stained with Giemsa solution. The number of colonies was counted with ImageJ software (National Institutes of Health).

### Annexin V Assay

Apoptosis was evaluated by flow cytometry with the Annexin V-FITC Apoptosis Kit (BioVision, Milpitas, CA, USA) according to the instructions of the manufacturer. Briefly, cells were harvested and stained with Annexin V-FITC and propidium iodide. Each sample was analyzed using the BD FACS Aria IIu (BD Biosciences, San Jose, CA, USA).

### RNA Isolation

Total mRNA was isolated with TRIzol Reagent (Invitrogen) for mouse samples and cell lines or with miRNeasy Mini Kit (Qiagen, Hilden, Germany) for human tissue samples, according to the respective manufacturer’s protocol. RNA quality was assessed with a NanoDrop ND-2000 spectrophotometer (Thermo Fisher Scientific).

### Real-time Quantitative RT-PCR Analysis

For quantitative analysis of mRNA, total RNA was reverse transcribed using the High Capacity RNA-to-cDNA Kit (Applied Biosystems). qRT-PCR was performed with a LightCycler 480 Real-Time PCR system (Roche Diagnostics, Mannheim, Germany) using the specific primers and LightCycler-DNA Master SYBR Green I (Roche Diagnostics) or with ABI Prism 7900HT Sequence Detection System (Applied Biosystems) using TaqMan Gene Expression Assays (Applied Biosystems) and TaqMan Universal PCR Master Mix (Applied Biosystems). The specifically designed primers and the product numbers of the TaqMan Gene Expression Assay were listed in Supplementary Table S5. Each gene expression value was normalized to the mRNA expression level of β-actin. Relative expression was quantified with the ΔΔCT method (Livak and Schmittgen, 2001).

### Clinical Tissue Samples

CRC samples were collected from 98 patients (Stage 0/I/II/III/IV: 5/28/29/28/8), of which 20 were early cancers. These patients underwent surgery between 2003 and 2010 at Osaka University Hospital. The Union for International Cancer Control classification was used for patient staging (Amin MB. American Joint Committee on Cancer). For transcriptome analysis, tissue samples were immediately frozen in RNAlater^TM^ (Ambion, Austin, TX, USA) and stored at -80°C until RNA extraction. For immunohistochemistry, tissue samples were fixed in 10% buffered formalin at 4°C overnight, processed through graded ethanol solutions, and embedded in paraffin.

### Ethics Approval

This study was approved by the institutional review board of our institution (Permission No. #15144). Written informed consent was obtained from all patients. The study protocol was in accordance with the Declaration of Helsinki, the Japanese Ethical Guidelines for Human Genome/Gene Analysis Research, and the Ethical Guidelines for Medical and Health Research Involving Human Subjects in Osaka University.

### Immunohistochemistry

Tissue sections of 4-μm thickness were prepared from paraffin-embedded blocks. H&E staining was performed for histological examination. Immunostaining was carried out with the Vectastain ABC Peroxidase Kit (Vector Laboratories, Burlingame, CA, USA), after antigen retrieval treatment in 10 mM citrate buffer (pH 6.0) at 95°C for 40 min. The slides were incubated overnight at 4°C with anti-human polyclonal antibody against MAF (#ab72584, Abcam), anti-mouse monoclonal antibody against p53 (#M7001, Dako, Glostrup, Denmark), anti-rabbit monoclonal antibody against SOX10 (#ab227680, Abcam), anti-rabbit polyclonal antibody against BCL6 (#PA5-27390, Invitrogen), anti-rabbit polyclonal antibody against S-100 (#422091, NICHIREI BIOSCIENCE Inc., Tokyo, Japan), anti-rabbit polyclonal antibody against CD45R (#14-0451-82, Invitrogen), and anti-rabbit monoclonal antibody against Ki-67 (#12202, Cell Signaling Technology, Danvers, MA, USA) at the following dilutions: anti-MAF antibody, 1:200; anti-p53, 1:50, anti-S100, 1:1; anti-SOX10, 1:100; anti-BCL6, 1:100; and anti-CD45R, 1:100.

Tissue sections of 8-μm thickness were prepared from OCT compound–embedded blocks. They were incubated overnight at 4°C with anti-mouse monoclonal rabbit antibody against Ki-67 (#12202, Cell Signaling Technology) at dilutions of 1:200. Counter nuclear staining was performed with a hematoxylin solution.

### Systemic Administration of miRNAs to *CPC;Apc* Mice

*Apc^min^* mice produce polyps mainly in the small intestine, whereas *CPC;Apc* mice produce colorectal tumors in which conditional knockout of the *Apc* gene was accomplished under the CDX2 promoter (∼9.5-kb 5’-flanking region from the human CDX2 gene), specifically acting at the mouse colorectum (Hinoi *et al*., 2007). Mice were treated with formulated miRNAs (miR-200c-3p, miR-302a/b/c/d-3p, and miR-369-3p/-5p) or negative control miRNA, three times a week for 8 weeks. miRNAs were injected via tail vein using sCA as a drug delivery system (Wu et al., 2015). The preparation of a sCA transfection mixture for *in vivo* use was previously described (Wu et al., 2015). Briefly, for a one-shot injection, 25 μg of each miRNA (the total amount of nucleic acid, 175 μg) or an equal amount of NC miRNA, and 350 μL of 1 M CaCl2 were mixed with 87.5 mL of serum-free bicarbonate–buffered inorganic solution (NaHCO3 44 mM, NaH2PO4 0.9 mM, CaCl2 1.8 mM, pH 7.5), and incubated at 37°C for 30 min. The solution was centrifuged at 12,000 rpm for 3 min and the pellet dissolved with saline containing 0.5% albumin. The products were sonicated (38 kHz, 80 W) in a water bath for 10 min to generate sCA plus 0.5% albumin, which was intravenously injected (approximately 70 μg per mouse) within 10 min. All miRNAs used in this study were purchased from Gene Design Inc. (Osaka, Japan; Supplementary Table S1).

At 15 weeks after birth, mice were sacrificed, and normal mucosa and polyps were collected, immediately frozen in RNAlater^TM^ (Ambion), and stored at -80°C until RNA extraction. All experiments were performed in strict accordance with the prescribed guidelines and protocols approved by the Committee on the Ethics of Animal Experiments of Osaka University (No. 30011026). Generation of colorectal polyps was monitored by a small-diameter rectosigmoid scope (Natsume Seisakusho, Tokyo, Japan).

### Microarray Analysis

Total RNA from mouse tissue samples was reverse transcribed with oligo-dT primers containing the T7 RNA polymerase promoter sequence. The resulting cDNA was subjected to *in vitro* transcription with T7 RNA polymerase for Cy3 labeling (CyDye; Amersham Pharmacia Biotech). Cy3-labeled cRNAs (600 ng) were hybridized onto Agilent Sure Print G3 Mouse GE 8×60K (G4852A). The signal intensity of Cy3 was calculated for every probe, and the results were analyzed with the Subio Basic Plug-in (v1.6; Subio Inc.), which allows for visualization of microarray data in the form of a heat map. The microarray raw data are available in the Gene Expression Omnibus (GEO; https://www.ncbi.nlm.nih.gov/geo/) database with accession code GSE92944.

### RNA Sequencing

We conducted RNA sequencing as previously described (Fukata et al., 2018). The library was prepared using a TruSeq Stranded mRNA Sample Prep Kit (Illumina, San Diego, CA, USA). Sequencing was performed using the Illumina HiSeq 2500 platform in 75-base single-end mode. Illumina Casava 1.8.2 software was used for base calling, and the sequenced reads were mapped to human reference genome sequences (hg19) using TopHat version 2.0.13 combined with Bowtie2 version 2.2.3 and SAMtools version 0.1.19. We calculated the fragments per kilobase of exon per million mapped fragments using Cuffnorm version 2.2.1. We identified a series of genes that were enhanced or reduced (tumor: 1.5 fold; normal mucosa: 1.3 fold) for further gene expression analysis. The raw data were deposited in the NCBI Gene Expression Omnibus database under GEO accession number GSE210970. We identified enhanced or suppressed pathways using Qiagen’s Ingenuity Pathway Analysis (IPA; Qiagen Redwood City, CA, USA; www.qiagen.com/ingenuity) with the default settings.

### Generation of c-MAF KO Mice

#### Mouse species

Jcl:ICR pseudopregnant female mice and C57BL/6JJcl cryopreserved zygotes were purchased from CLEA Japan Inc. (Tokyo, Japan). All animals were housed and maintained under conditions of 50% humidity and a 12:12-h light:dark cycle. They were fed a standard pellet diet (MF, Oriental Yeast Co., Tokyo, Japan) and tap water *ad libitum*. The Osaka University Animal Experiment Committee approved all animal experiments.

#### Preparation of Cas9 and gRNA

The following reagents were purchased: Cas9 protein, Alt-R® S.p. Cas9 Nuclease 3NLS (Integrated DNA Technologies, Inc. USA); guide RNA (gRNA), and GeneArt Precision gRNA Synthesis Kit (Thermo Fisher Scientific). To design the gRNA sequence (5’-CAGGAGGATGGCTTCAGAAC-3’), we used a software tool (http://crispr.mit.edu/) to predict unique target sites throughout the mouse genome.

#### Electroporation into mouse embryos

Pronuclear-stage mouse embryos were prepared by thawing frozen embryos in KSOM medium (ARK Resource, Kumamoto, Japan). For electroporation, 150 embryos at 1 h after thawing were placed into a chamber with 40 µL of serum-free medium (Opti-MEM, Thermo Fisher Scientific) containing 100 ng/µL Cas9 protein and 200 ng/µL gRNA. They were electroporated with a 5-mm gap electrode (CUY505P5 or CUY520P5 Nepa Gene, Chiba, Japan) in a NEPA21 Super Electroporator (Nepa Gene, Chiba, Japan). The poring pulses for the electroporation were voltage 225 V, pulse width 1 ms, pulse interval 50 ms, and number of pulses 4. The first and second transfer pulses were voltage 20 V, pulse width 50 ms, pulse interval 50 ms, and number of pulses 5. Mouse embryos that developed to the two-cell stage after the introduction of Cas9 and gRNA were transferred into the oviducts of female surrogates anesthetized with sevoflurane (Mylan Pfizer Japan Inc.). Male and female mice with c-MAF heterogeneous KO (MAF+/-) were mated so that homogeneous *c-MAF* KO (MAF-/-) c-MAF and heterogeneous KO (MAF+/-) mice were produced.

#### Genotyping analysis

Genomic DNA was extracted from the tail tip using the KAPA Express Extract DNA Extraction Kit (Kapa Biosystems, London, UK) and Animal Tissue Direct PCR Amplification Kit (with TL) (FineGene, Shanghai, CN). For PCR and sequence analysis, we used primers that amplified the targeted region. PCR was performed under the following conditions: 1 cycle of 94°C for 1 min; 30 cycles of 98°C for 10 s, 60°C for 15 s, and 68°C for 30 s; and 1 cycle of 72°C for 3 min. The PCR products were sequenced immediately or after purification using the Mini-Gel extraction kit (One-Step) (FineGene) with BigDye Terminator v3.1 cycle sequencing mix and the standard protocol for an Applied Biosystems 3130 DNA Sequencer (Life Technologies).

### Examination of Tumorigenesis in Mice

#### Spontaneous carcinogenesis

c-MAF KO mice (n = 187) and wild-type mice (n = 37) were observed for 24 months for tumor generation. Tumors were histologically examined with H&E staining and the specific immunostaining.

#### Chemical carcinogenesis

AOM and DSS treatment was used to produce CRCs, as previously reported (Parang et al., 2016). On day 100 after administration of AOM, all mice were examined by an animal endoscope for tumor formation in the colorectum (Natsume Seisakusho). On 136 ± 8 day, mice were sacrificed, and the colorectum was removed. After gross observation, normal mucosa and tumors of diameter >1 mm were collected and subjected to histological examination and RNA sequencing.

### Statistical Analysis

All data are expressed as mean ± standard deviation, or the median and interquartile range (IQR). Statistical differences were analyzed using Student’s *t* test for continuous variables and the Chi-squared test for non-continuous data. Survival curves were developed with the Kaplan–Meier method and compared using the log-rank test. A Cox proportional hazard regression model was used to estimate HRs and 95% CIs. All statistical analyses were conducted with JMP ver. 14.0 (SAS Institute, Inc., Cary, NC, USA). All *P* values < 0.05 were considered to indicate statistical significance.

**Supplementary Fig. S1. Heat map analysis of normal mucosa and polyps in *CpC;Apc* mice treated with the miR cocktail (miRs: 200c, 302a-d, 369) or with negative control (NC) miRNA.** A subset of normal mucosa and polyps with RIN (RNA integrity number) >6.3 were subject to microarray analysis (miRs-Normal, n = 2; NC miR-Normal, n = 2; NC miR-Polyp, n = 7). A total of 15 genes were highly expressed by polyps, and 53 genes showed stepwise downregulation from the miR-treated normal mucosa to NC-treated normal mucosa to NC-treated polyps.

**Supplementary Fig. S2. Database survey of c-MAF mRNA expression in human cancers.** (A) c-MAF mRNA level decreased in colon and rectum. The public database ONCOMINE^TM^ indicated that c-MAF mRNA expression was decreased in adenocarcinoma of colon and rectum compared with normal mucosa (Rhodes et al., 2004). (B) Comparison of c-MAF mRNA expression between tumor and normal tissues in a series of human malignancies (http://firebrowse.org/). (C) Many human malignancies, including colon (COAD) and rectal cancer (READ) or both (COADREAD) expressed lower levels of c-MAF mRNA as compared with their normal counterparts.

**Supplementary Fig. S3. Genotyping of wild-type and *c-MAF* KO mice.** Genomic DNA was extracted from the tail tip, the target region was amplified by PCR, and Sanger sequencing analysis was performed. Red arrowheads indicate the deletion start site viewed from the 3’ side and deleted sequences are indicated by the dotted line (Supplementary Fig. S3A). To clarify deletion sites in comparison with the wild-type sequence, the corresponding sequences were marked in blue, green, and red (Supplementary Fig. S4A). Black arrows on the right indicate read direction; note that heterogeneous KO region displays overlapped peaks on the 5’ side from the red arrowhead while the homogeneous KO region showed a single peak (Supplementary Fig. S3B).

**Supplementary Fig. S4. Spontaneous carcinogenesis in c-MAF KO mice.** (A) One mouse (8-nucleotide deletion/c-MAF hetero) had a prolapse of the bowel. Gross examination of the removed colorectum revealed that it was a rectal tumor. Histologically, it was a well- to moderately differentiated adenocarcinoma (scale bar: left; 500 μm, right; 100 μm). (B) In one mouse (one-nucleotide deletion/c-MAF hetero), a pale white tumor was observed on the chest wall (scale bar: 1 cm). By H&E and immunohistochemical staining (S100-positive, SOX10-negative), the tumor was diagnosed as a schwannoma (scale bar: 100 μm). (C) In one mouse (one-nucleotide deletion/c-MAF hetero), a tumor that tightly adhered to the small intestine was observed (scale bar: 1 cm). Through histological examination and immunohistochemical analysis, it was weakly BCL6 positive and CD45R positive and diagnosed as Burkitt’s lymphoma (scale bar: 100 μm).

**Supplementary Fig. S5. colitis-associated tumor formations in c-MAF knockout mice.** (A) Experiments of chemical carcinogenesis were performed. Azoxymethane (AOM) was intraperitoneally injected on day 0, and 2% DSS was administered in drinking water on days 7–11 and 28–32. (B) Tumors were distributed predominantly in the rectum and distal colon. (C) Prolapse of a rectal cancer. (I) On day 100, one mouse had a prolapse from the anus, indicated by the yellow arrowhead in (a). (II) Endoscopic examination confirmed it was rectal tumor (yellow arrowhead in a) and revealed that multiple tumors existed. (III) Gross findings of the removed colon and rectum. A rectal tumor indicated by the yellow arrowhead (a) and another two polyps (indicated by arrowheads) were noted. (IV) H&E staining of the rectal tumor. Histological diagnosis was well- to moderately differentiated adenocarcinoma. Scale bar: 500 μm. (V) Magnification view of the rectangle area. Scale bar: 100 μm.

**Supplementary Fig. S6. RNA sequencing of colorectal tissue samples of wild-type and *c-MAF KO* mice after treatment with AOM/DSS.** (A) The heat map of upregulated or downregulated gene expression in normal colonic mucosa between *c-MAF* KO mice and wild-type (WT) mice. (B) Heat map of gene expression in tumors between *c-MAF* KO mice and WT mice.

**Supplementary Fig. S7. Ingenuity Pathway Analysis of downstream disease and function in tumors of *c-MAF* KO mice**. A number of cancer-related categories were activated in *c-MAF* KO tumors compared with wild-type tumors.

**Supplementary Fig. S8. Effect on miR-155 induction of transduction of p53 mutation into p53-null HCT116 cells.** (A) The mutated p53 plasmids (R175H, R248W) were transfected into p53-null HCT116 cells to produce p53 protein. The HT29 cells harboring the mutated p53 gene served as a positive control. Actin bands served as a loading control. EV: empty vector. (B) c-MAF expression was reduced in mutated p53-transfected cells. (C) qRT-PCR revealed that miR-155 levels significantly increased in mutated p53-transfected cells compared with vector control; \**P* < 0.05. (D) Expression of c-MAF was decreased in the miR-155–transfected cells.

**Supplementary Fig. S9. A candidate mechanism for how miRs-200c, -302a-d, and - 369 induce c-MAF.** (A) Ingenuity Pathway Analysis was used to create a gene relationship diagram centered on c-MAF. The diagram revealed that the transcription factor GATA3 positively regulates MAF expression, as previously reported (Naito et al., 2011). RNA sequencing of MRC5 cell contents at 48 h after transfection by miRs- 200c, 302a-d, and 369s indicated that GATA3 mRNA expression had a 3.07-fold increase as compared with basal level in parental MRC5 cells. (B) In human MRC5 fibroblasts and HBEC3-KT human bronchial epithelial cells, treatment with the miRNAs increased GATA3 and c-MAF mRNA expression (\*\**P* < 0.01).

## Notes

### Competing Interest Statement

The authors have declared no competing interest.

