## supplementary figure for "Paradoxical tumor suppressive role of the *musculoaponeurotic fibrosarcoma* gene in colorectal cancer"

#### Supplementary Fig. S1

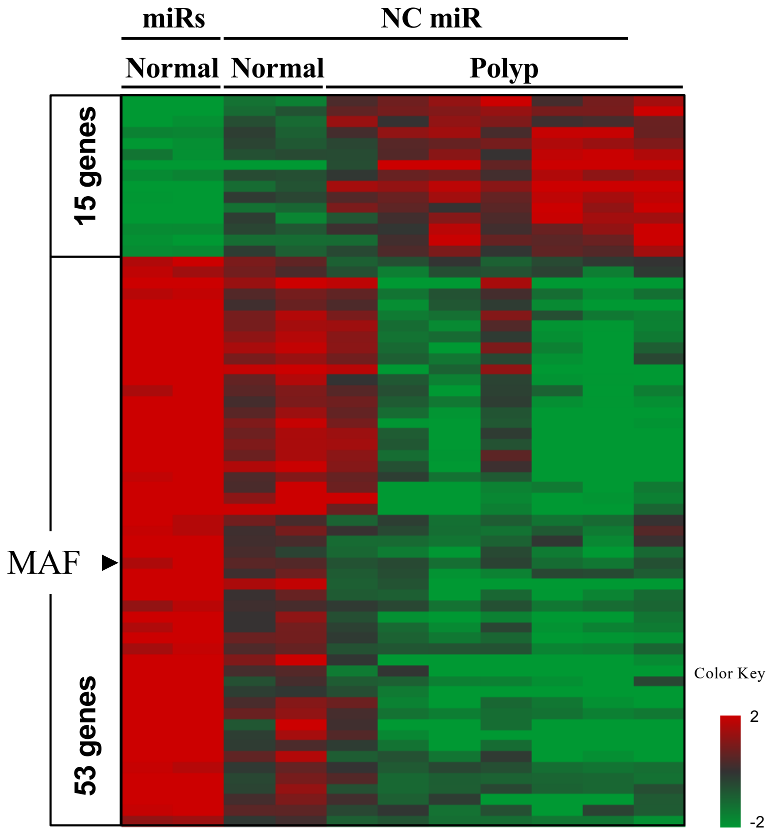

**Supplementary Fig. S2**

A

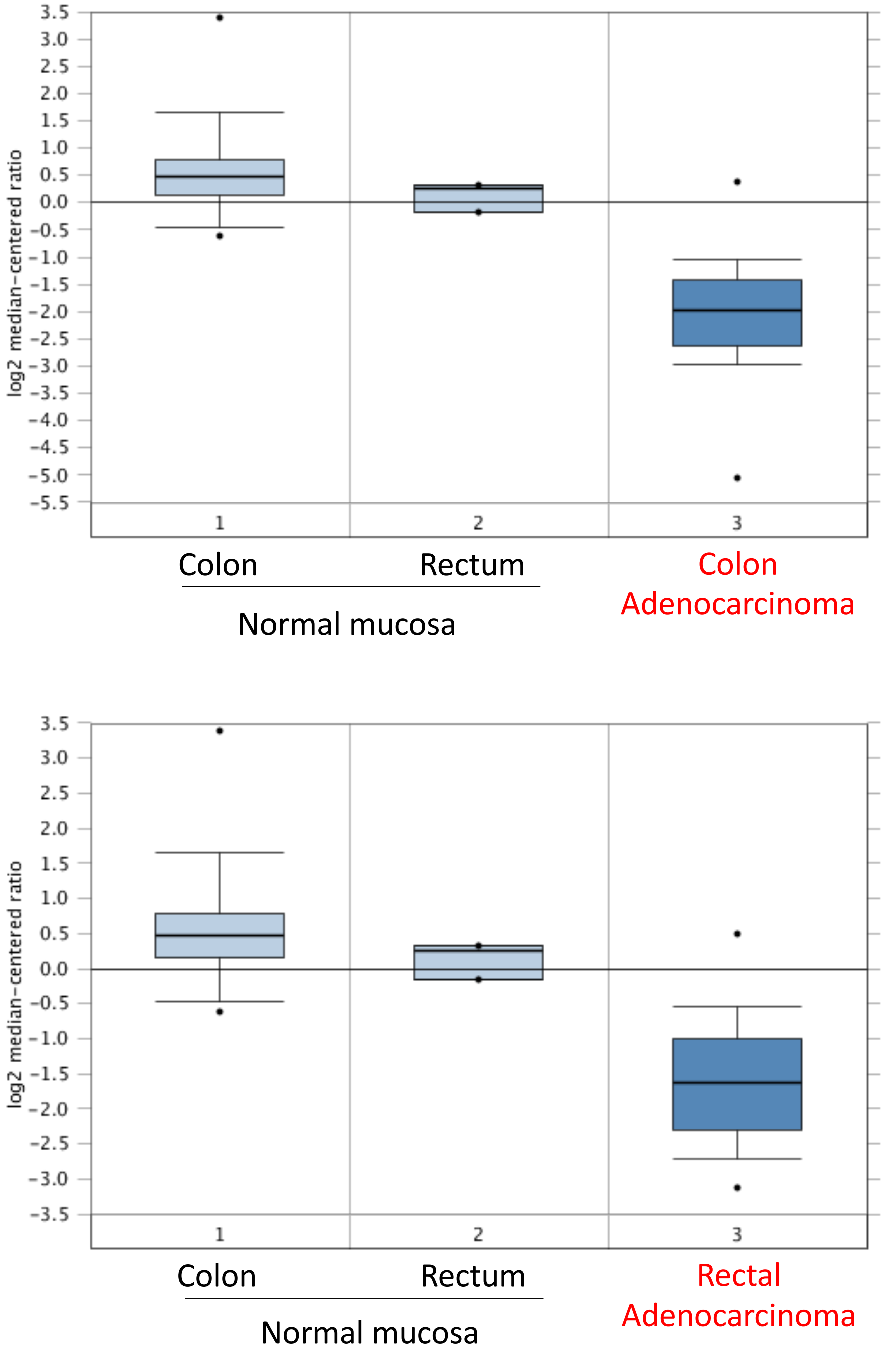

**Supplementary Fig. S2**

B

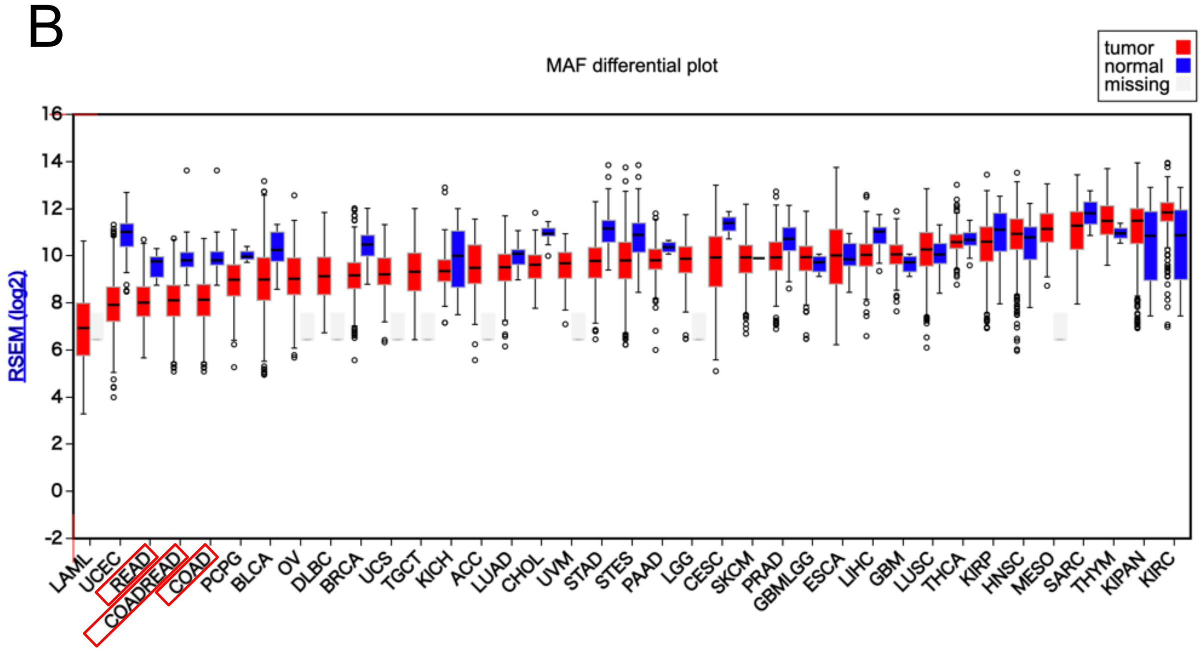


| Abbreviation List | |
|---|---|
| LAML | Acute myeloid leukemia |
| UCEC | Uterine corpus endometrial carcinoma |
| READ | Rectum adenocarcinoma |
| COADREAD | Colorectal adenocarcinoma |
| COAD | Colon adenocarcinoma |
| PCPG | Pheochromocytoma and paraganglioma |
| BLCA | Bladder urothelial carcinoma |
| OV | Ovarian serous cystadenocarcinoma |
| DLBC | Lymphoid neoplasm diffuse large b-cell lymphoma |
| BRCA | Breast invasive carcinoma |
| UCS | Uterine carcinosarcoma |
| TGCT | Testicular germ cell tumors |
| KICH | Kidney chromophobe |
| ACC | Adrenocortical carcinoma |
| LUAD | Lung adenocarcinoma |
| CHOL | Cholangiocarcinoma |
| UVM | Uveal melanoma |
| STAD | Stomach adenocarcinoma |
| STES | Stomach and esophageal carcinoma |
| PAAD | Pancreatic adenocarcinoma |
| LGG | Brain lower grade glioma |
| CESC | Cervical squamous cell carcinoma and endocervical adenocarcinoma |
| SKCM | Skin cutaneous melanoma |
| PRAD | Prostate adenocarcinoma |
| GBMLGG | Glioma and low-grade glioma |
| ESCA | Esophageal carcinoma |
| LIHC | Liver hepatocellular carcinoma |
| THCA | Thyroid carcinoma |
| KIRP | Kidney renal clear cell carcinoma |
| HNSC | Head and neck squamous cell carcinoma |
| MESO | Mesothelioma |
| SARC | Sarcoma |
| THYM | Thymoma |
| KIPAN | Pan-kidney cohort |
| KIRC | Kidney renal clear cell carcinoma |

**Supplementary Fig. S2**

C

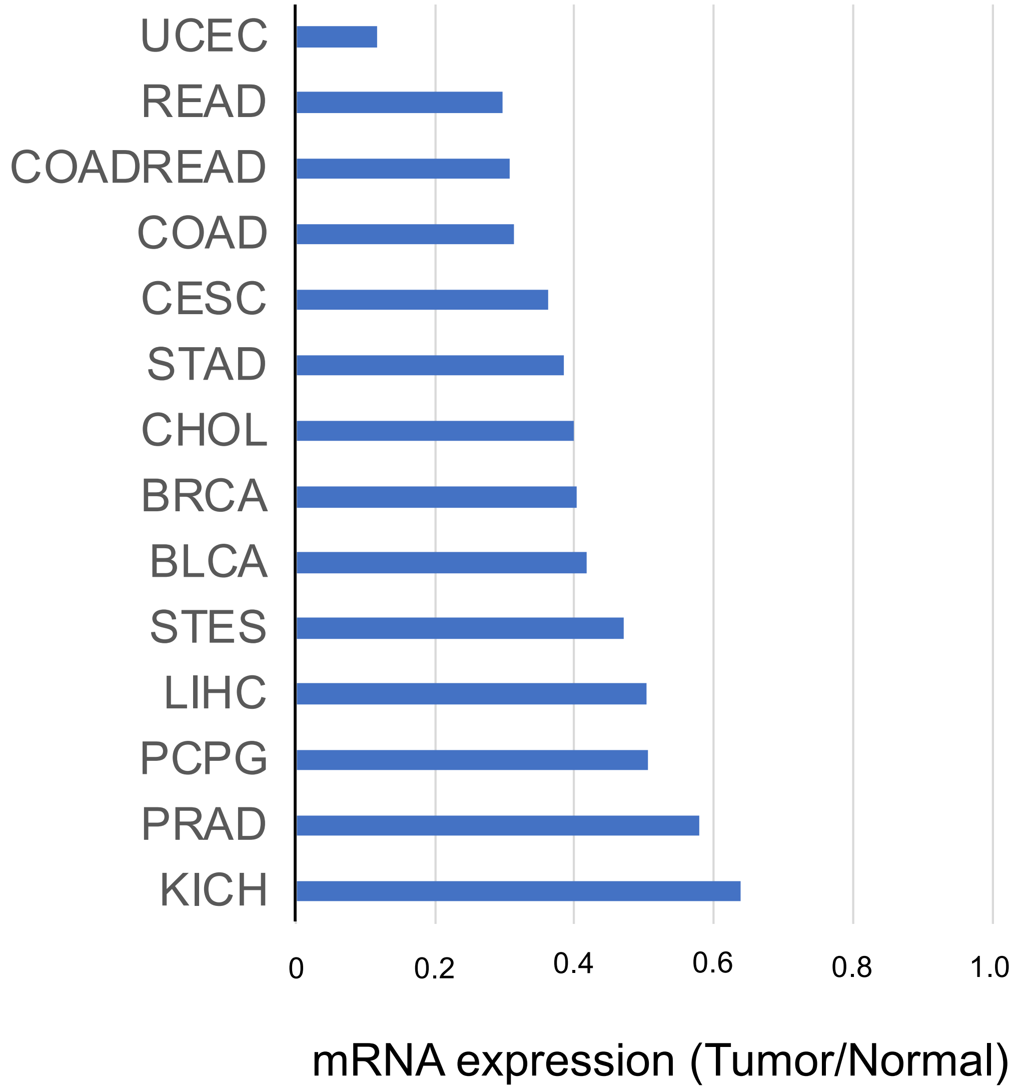

#### Supplementary Fig. S3A

| Type | | Sangar sequence (5’ to 3') |
|---|---|---|
| Wild type | | GCTGGCCACCCAGCACAGCCCGCTGGCCCCTGTCCTGCAGCCCATCTGGCGGAGCGGCGGC<br>GGCGGTGGCGGCGGAGGCAGGAGGATGGCTTCAGAACTGGCAATGAACAATTCCGACCTGC<br>CCACCAGTCCCCTGGCCATGGAATATGTTAATGACTTCGATCTGATGAAGTTTGAAGTGAAAAA<br>GGAACCGGTGGAGACCGACCGCATCATCAGCCAGTGCGGCCGTCTCATCGCCGGGGGCTCG<br>CTGTCCTCCACCCCCATGAGCACGCCCTGCAGCTCGGTGCCCCCNTCNNNCANGNTNNNNNNN<br>NNNNNNNNNN |
| 1 nucleotide deletion | homo | NNNTGTGCACGTTCGAGCTTTCGGGCCACAGCCACCGCCGCGCAAGCTAGAAGCGCCCTAGC<br>CCGGCAAGCTGGCTCACCCGCTGGCCACCCAGCACAGCCCGCTGGCCCCTCTCCTGCAGCCC<br>ATCTGGCGGAGCGGCGGCGGCGGTGGCGGCGGAGGCAGGAGGATGGCTTCA-AACTGGCAA<br>TGAACAATTCCGACCTGCCCACCAGTCCCCTGGCCATGGAATATGTTAATGACTTCGATCTGAT<br>GAAGTTTGAAGTGAAAAAGGAACCGGTGGAGACCGACCGCATCATCAGCCAGTGCGGCCGTCT<br>CATCGCCGGGGGCTNNNNNNCCTCCACCCCCATGAGCACGCCCTGCAGCTCGGTGCCCCCNT<br>CCCCCAGCNNCTNGGNNNNNNNNNNN |
| | hetero | NNNNNNNNNNNNNTNNNNNNNNTNGGGNCNNNNNCCNCNNNNNNNCCANNNNNNANNNNCNNNNN<br>CCNGNNANNNNGNNNNNCCNNNNGNCCNNCNNNNNNNNNCCNNNGGNCCCTTTTNNNNNNNCC<br>NTTNNGNNGNNNNGNNGNGGNNGNNGNNGNNGNNGCCNGNNGNNNGNNTTNNAACTGGCAA<br>TGAACAATTCCGACCTGCCCACCAGTCCCCTGGCCATGGAATATGTTAATGACTTCGATCTGAT<br>GAAGTTTGAAGTGAAAAAGGAACCGGTGGAGACCGACCGCATCATCAGCCAGTGCGGCCGTCT<br>CATCGCCGGGGGCTCGCTGTCCTCCACCCCCATGAGCACGCCCTGCAGCTCGGTGCCCCCNT<br>CCCCCAGCTCTNGGCNCCNNNNNNN |
| 8 nucleotide deletion | homo | NGTGTGCACGTTCGAGCTTTCGGGCCACAGCCACCGCCGCGCAAGCTAGAAGCGCCCTAGCC<br>CGGCAAGCTGGCTCACCCGCTGGCCACCCAGCACAGCCCGCTGGCCCCTCTCCTGCAGCCCA<br>TCTGGCGGAGCGGCGGCGGCGGTGGCGGCGGAGGCAGGAGGATGGCT--------GGCAATGAA<br>CAATTCCGACCTGCCCACCAGTCCCCTGGCCATGGAATATGTTAATGACTTCGATCTGATGAAG<br>TTTGAAGTGAAAAAGGAACCGGTGGAGACCGACCGCATCATCAGCCAGTGCGGCCGTCTCATC<br>GCCGGGGGCNNNNNNTCCTCCACCCCCATGAGCACGCCCTGCAGCTCGGTGCCCCCNTCCCC<br>CAGCTCTNGGCNNNCNNNN |
| | hetero | TGTGTGCNTGTGTGCACNTTCGAGCCTCAGGCCCCCACGCGCAAGCGAGCAAGCTAGAAGCC<br>CGGCAAGCTGGCTAGCTGGCTCACCCGCCGGCCCACCAGCACAGCCCGTTTCCCCCTGTCCA<br>TCAGGCCGTGCGGCGGAGCGGGGGGGGCGGGGGCCAGGGGGACAGGATGAGGGCTGGCA<br>ATGAACAATTCCGACCTGCCCACCAGTCCCCTGGCCATGGAATATGTTAATGACTTCGATCTGA<br>TGAAGTTTGAAGTGAAAAAGGAACCGGTGGAGACCGACCGCATCATCAGCCAGTGCGGCCGTC<br>TCATCGCCGGGGGCTCGCTGTCCTCCACCCCCNATGAGCNCGCCCTGCTGCTCGGTGCCCCC<br>GTCCCCCCAGCTCTCGGCNCCCNNNC |

**Fig. S3B**

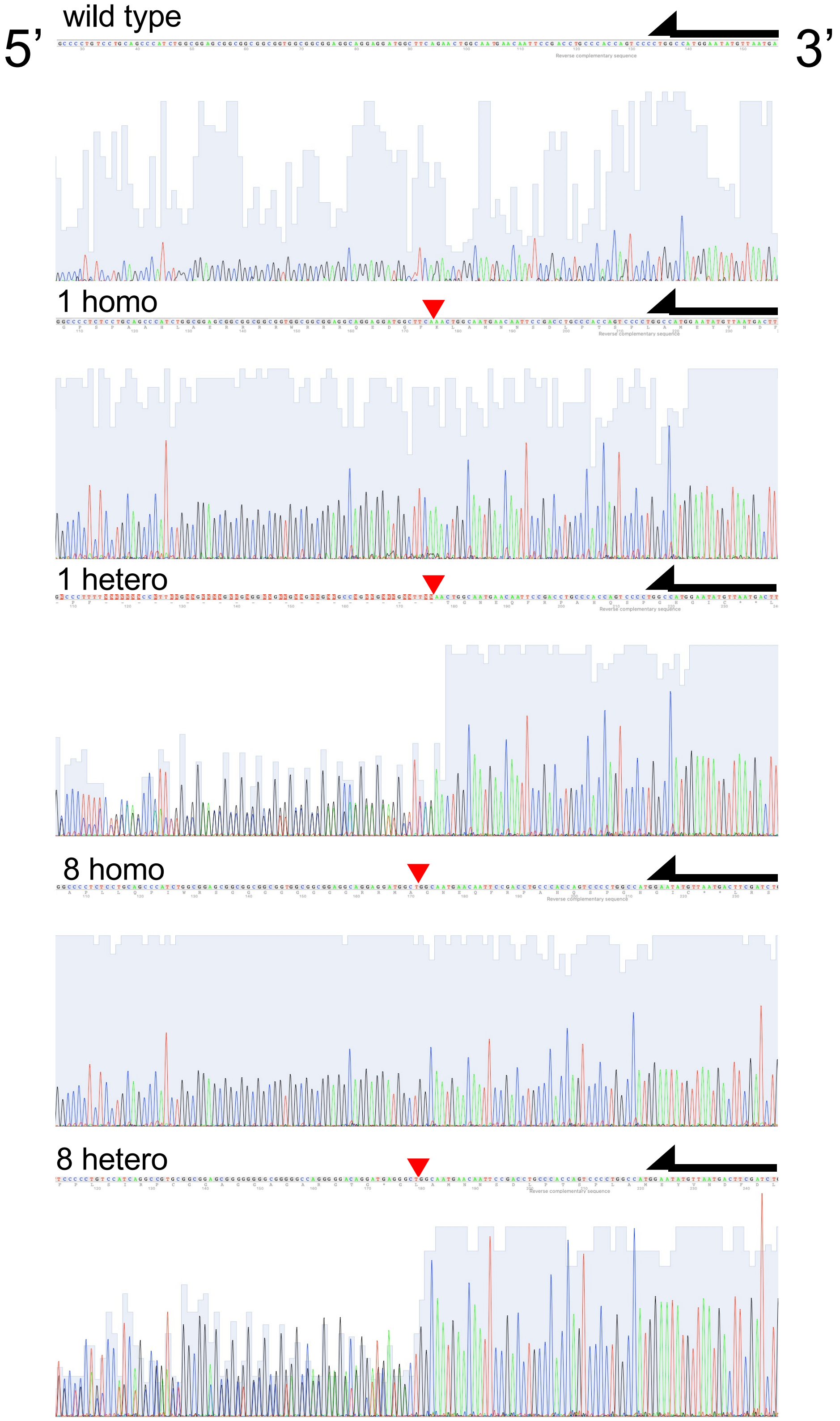

#### Supplementary Fig. S4

A

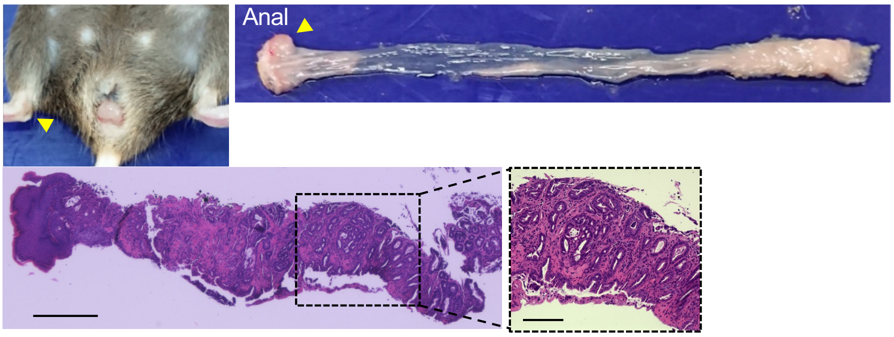


B

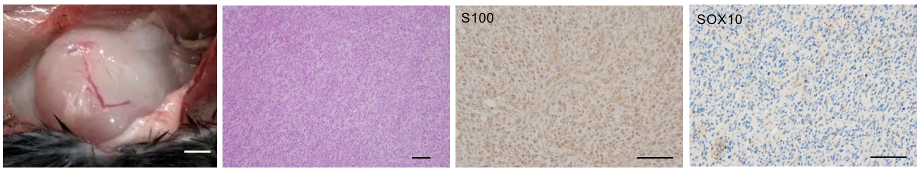


C

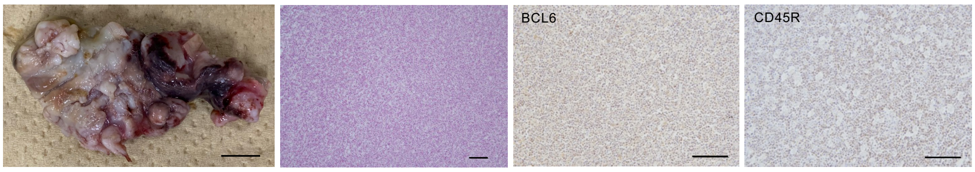

#### Supplementary Fig. S5

A

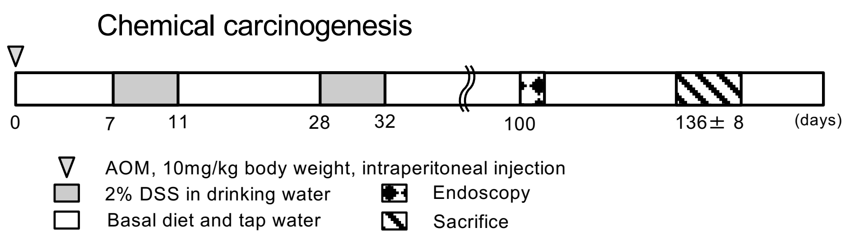


B

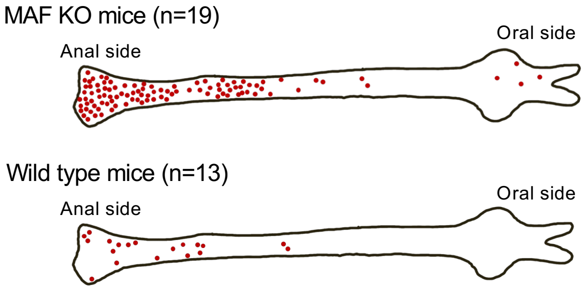


C

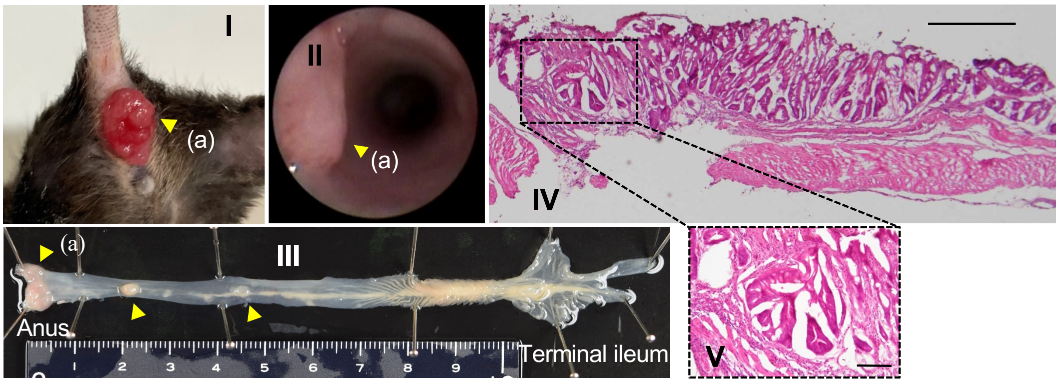

#### Supplementary Fig. S6

**A**

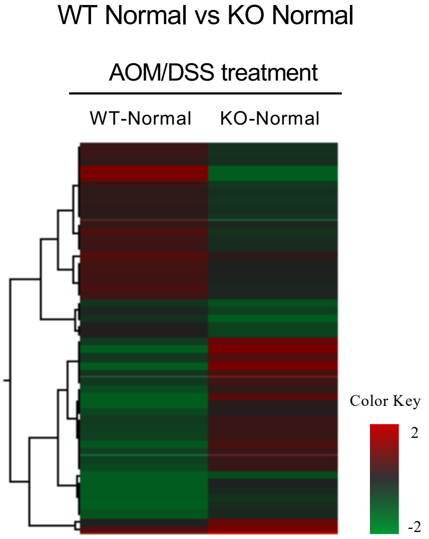


WT-Normal/KO-Normal
Up & Down Top 25

| ID | Fold Change | *P* value |
|---|---|---|
| Xist | 94.870 | 0.009 |
| Hba-a2 | 58.735 | 0.043 |
| Eif3j1 | 33.139 | 0.040 |
| Adig | 6.726 | 0.001 |
| Itln1 | 6.578 | 0.033 |
| Pnliprp2 | 6.041 | 0.013 |
| Cpa2 | 5.731 | 0.042 |
| Clps | 3.942 | 0.008 |
| Gsdmc | 3.844 | 0.004 |
| Duoxa1 | 3.763 | 0.003 |
| Car3 | 3.693 | 0.027 |
| S100a9 | 3.410 | 0.007 |
| Tpsb2 | 3.379 | 0.007 |
| Cryba4 | 3.368 | 0.010 |
| Tuba8 | 3.160 | 0.004 |
| Clca1 | 3.150 | 0.012 |
| Gm13212 | 3.081 | 0.003 |
| Ly6c2 | 3.062 | 0.021 |
| Adgre4 | 3.029 | 0.028 |
| Ear2 | 2.992 | 0.037 |
| Tg | 2.962 | 0.040 |
| C130026I21Rik | 2.879 | 0.003 |
| Chst13 | 2.806 | 0.015 |
| Gsdmcl-ps | 2.648 | 0.021 |
| Fpr2 | 2.617 | 0.007 |
| Eif2s3y | 0.013 | 0.012 |
| Ddx3y | 0.020 | 0.016 |
| C330022C24Rik | 0.331 | 0.024 |
| Gm10768 | 0.367 | 0.030 |
| Neat1 | 0.400 | 0.029 |
| Fam19a3 | 0.418 | 0.013 |
| Ddx60 | 0.446 | 0.004 |
| Myom3 | 0.446 | 0.040 |
| B430010I23Rik | 0.447 | 0.029 |
| Oasl2 | 0.451 | 0.028 |
| Apoo-ps | 0.453 | 0.038 |
| Ccdc116 | 0.460 | 0.027 |
| Gm3558 | 0.480 | 0.009 |
| Apol9a | 0.498 | 0.042 |
| 2410018L13Rik | 0.527 | 0.032 |
| Gm10069 | 0.535 | 0.031 |
| Rhov | 0.536 | 0.020 |
| Ido1 | 0.543 | 0.003 |
| 4933430I17Rik | 0.555 | 0.010 |
| Vsig10l | 0.560 | 0.049 |
| BC051142 | 0.561 | 0.020 |
| 5031434O11Rik | 0.561 | 0.026 |
| Npb | 0.571 | 0.050 |
| Slc4a4 | 0.572 | 0.045 |
| Ttc16 | 0.573 | 0.036 |

### Supplementary Fig. S6

**B**

WT Tumor vs KO Tumor

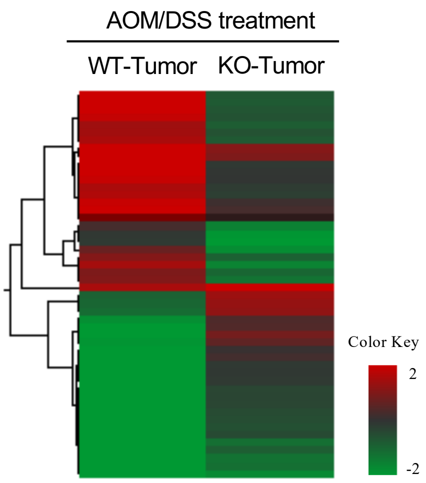


WT-Tumor/KO-Tumor
Up & Down Top 25

| ID | Fold Change | *P* value |
|---|---|---|
| 0610040F04Rik | 11.850 | 0.003 |
| Ctxn3 | 10.868 | 0.023 |
| Mab21l2 | 7.133 | 0.018 |
| Angptl7 | 6.820 | 0.029 |
| Vip | 6.722 | 0.031 |
| B3gnt6 | 5.998 | 0.017 |
| Hand2 | 5.646 | 0.004 |
| Dmrta2 | 5.645 | 0.002 |
| Prph | 5.487 | 0.007 |
| Gdf10 | 5.475 | 0.038 |
| Sst | 5.367 | 0.004 |
| Matn4 | 5.241 | 0.005 |
| H19 | 5.115 | 0.019 |
| Fzd9 | 5.094 | 0.001 |
| Mfap4 | 5.066 | 0.001 |
| Fmod | 4.873 | 0.005 |
| Upp2 | 4.791 | 0.043 |
| Snrpn | 4.750 | 0.003 |
| Kcnh3 | 4.702 | 0.007 |
| Calb2 | 4.572 | 0.006 |
| Acaa1b | 4.470 | 0.007 |
| Tmem59l | 4.421 | < 0.001 |
| Nov | 4.288 | 0.007 |
| Pou2f3 | 4.261 | 0.001 |
| Col8a2 | 4.197 | 0.033 |
| Hoxd11 | 0.049 | 0.013 |
| Hoxd10 | 0.073 | 0.017 |
| Hoxd13 | 0.080 | 0.027 |
| Mettl7a2 | 0.119 | 0.007 |
| Atp12a | 0.148 | 0.004 |
| Cps1 | 0.178 | 0.022 |
| Cyp2d12 | 0.184 | 0.001 |
| Lix1 | 0.193 | 0.025 |
| Gzmk | 0.196 | 0.012 |
| Efhb | 0.204 | 0.001 |
| S100g | 0.206 | 0.001 |
| Zp2 | 0.212 | 0.038 |
| Klhl14 | 0.236 | 0.008 |
| Adh6a | 0.239 | 0.031 |
| Abcg5 | 0.242 | 0.008 |
| Slc15a1 | 0.256 | < 0.001 |
| Lrrn3 | 0.257 | 0.003 |
| Timp4 | 0.258 | 0.022 |
| Cyp2c68 | 0.265 | 0.002 |
| Trpv3 | 0.274 | < 0.001 |
| Cyp2d9 | 0.277 | 0.002 |
| Scnn1g | 0.291 | 0.014 |
| Tgm3 | 0.291 | 0.038 |
| Ddx60 | 0.293 | 0.006 |
| Abcg8 | 0.303 | 0.031 |

#### Supplementary Fig. S7

#### KO tumors vs Wild type tumors

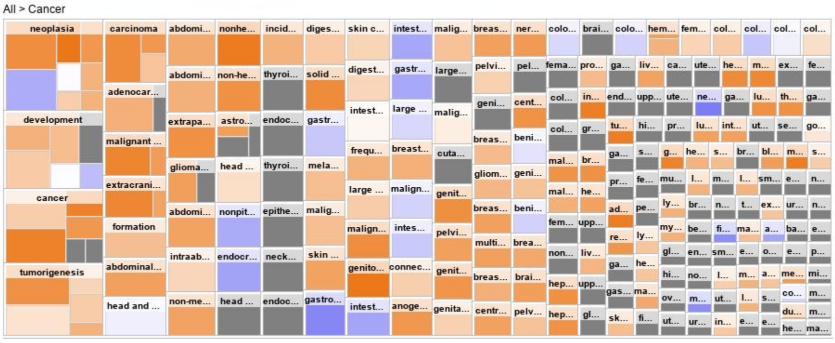


Colored by : z—score

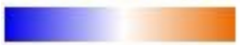

#### Supplementary Fig. S8

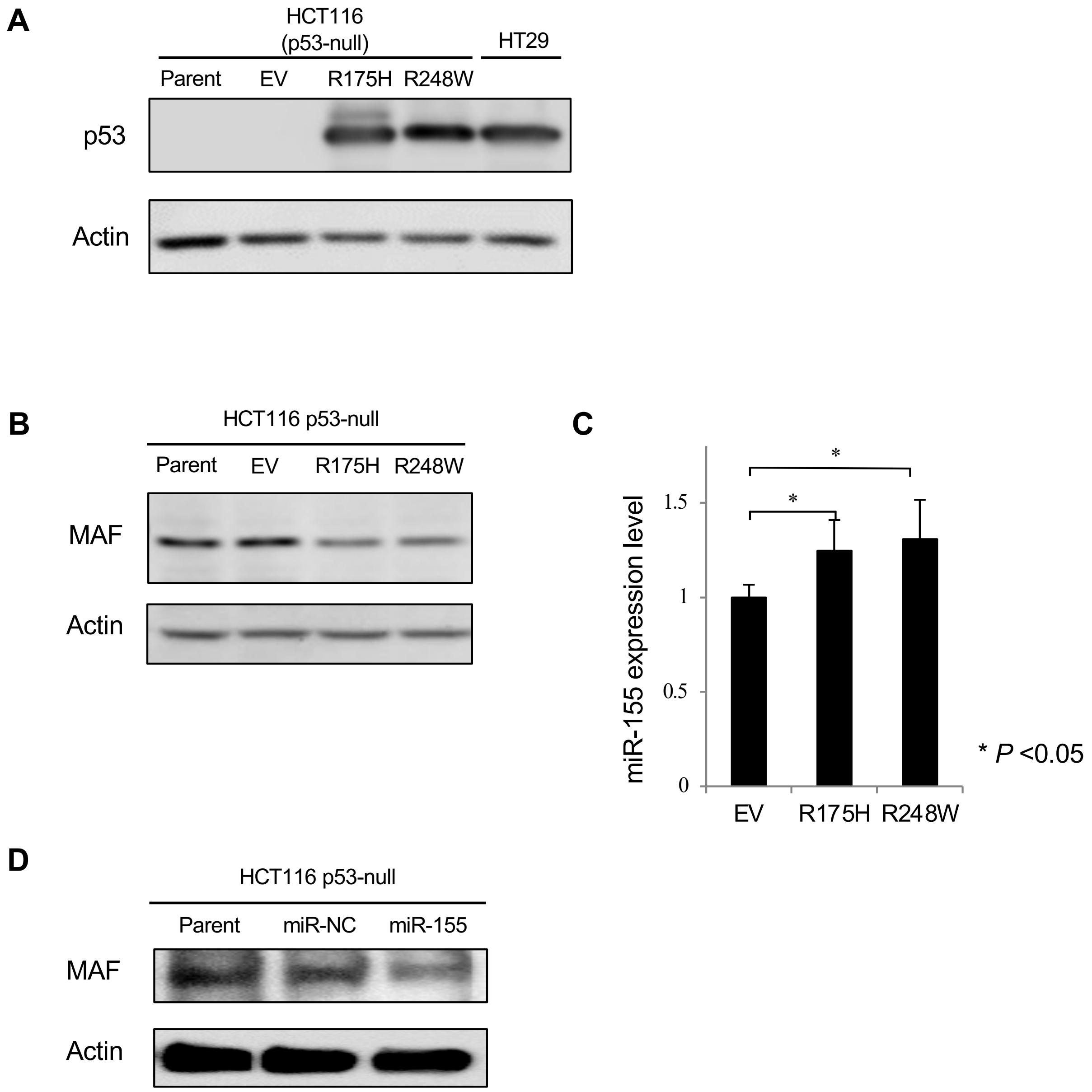

#### Supplementary Fig. S9

**A**

**RNA seq**

| ID | *P* value | Fold Change |
|---|---|---|
| GATA3 | 0.04 | 3.07 |

(miR/Parent)

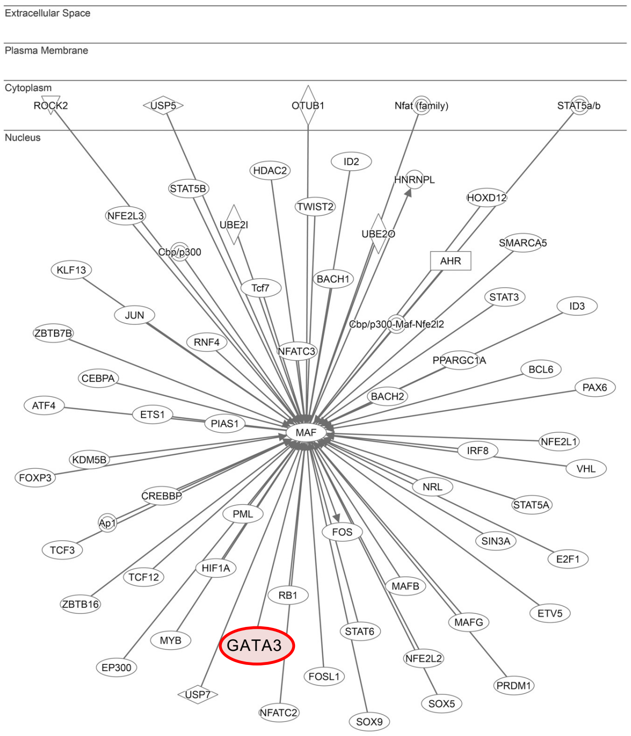


**B**

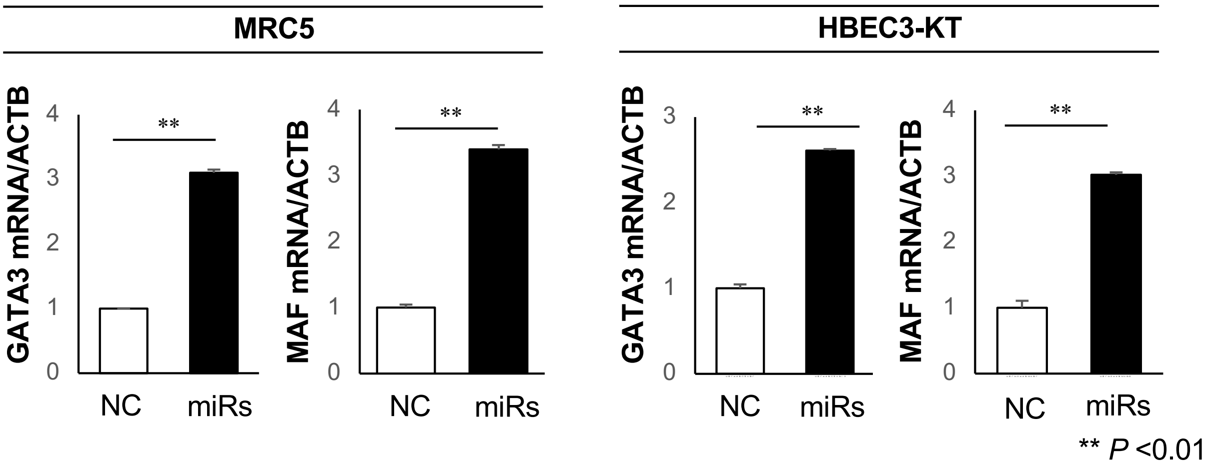
