## supplementary table for "Paradoxical tumor suppressive role of the *musculoaponeurotic fibrosarcoma* gene in colorectal cancer"

**Supplementary Table. S1. Sequences of miRNAs in human, mouse, and rat species, sense (5’ to 3’)**

| | hsa (human) | mmu (mouse) | rno (rat) |
|---|---|---|---|
| miR-200c-3p | UAAUACUGCCGGGUAAUGAUGGA | UAAUACUGCCGGGUAAUGAUGGA | UAAUACUGCCGGGUAAUGAUG |
| miR-302a-3p | UAAGUGCUUCCAUGUUUUGGUGA | UAAGUGCUUCCAUGUUUUGGUGA | N/A |
| miR-302b-3p | UAAGUGCUUCCAUGUUUUAGUAG | UAAGUGCUUCCAUGUUUUAGUAG | N/A |
| miR-302c-3p | UAAGUGCUUCCAUGUUUCAGUGG | AAGUGCUUCCAUGUUUCAGUGG | N/A |
| miR-302d-3p | UAAGUGCUUCCAUGUUUGAGUGU | UAAGUGCUUCCAUGUUUGAGUGU | N/A |
| miR-369-3p | AAUAAUACAUGGUUGAUCUUU | AAUAAUACAUGGUUGAUCUUU | AAUAAUACAUGGUUGAUCUUU |
| miR-369-5p | AGAUCGACCGUGUUAUAUUCGC | AGAUCGACCGUGUUAUAUUCGC | AGAUCGACCGUGUUAUAUUCGC |
| miR-155 | UUAAUGCUAAUCGUGAUAGGGGU | | |
| Negative control | UAAAUGUACUGCGCGUGGAGAGGAA | UCACAACCUCCUAGAAAGAGUAGA | |

**Supplementary Table. S2. Lists of activated molecules in tumors of *c-MAF* KO mice compared with those in wild-type mice after treatment with AOM/DSS.**

| Probe ID | | Gene Symbol | Description |
|---|---|---|---|
| 15 genes: Upregulation | A_55_P2032678 | Lox | Lysyl Oxidase (Lox) |
| | A_55_P2085974 | Igf1 | Insulin-Like Growth Factor 1 (Igf1), Transcript Variant 1 |
| | A_51_P409349 | Trim29 | Tripartite Motif-Containing 29 (Trim29) |
| | A_51_P264769 | Dmp1 | Dentin Matrix Protein 1 (Dmp1) |
| | A_55_P2436438 | Dusp22 | Dual Specificity Phosphatase 22 (Dusp22), Transcript Variant 1 |
| | A_55_P1998066 | Tmem121 | Transmembrane Protein 121 (Tmem121) |
| | A_51_P171616 | Wnt10a | Wingless Related Mmtv Integration Site 10A (Wnt10A) |
| | A_51_P363338 | Prkd1 | Protein Kinase D1 (Prkd1) |
| | A_55_P2000409 | Rab44 | Rab44, Member Ras Oncogene Family (Rab44) |
| | A_55_P2184796 | Pcdhb18 | Protocadherin Beta 18 (Pcdhb18) |
| | A_51_P436652 | Ccl7 | Chemokine (C-C Motif) Ligand 7 (Ccl7) |
| | A_51_P428372 | Ppbp | Pro-Platelet Basic Protein (Ppbp) |
| | A_55_P2145459 | 4930478M13Rik | Adult Male Testis Cdna, Riken Full-Length Enriched Library, Clone:4930478M13 Product:Unclassifiable, Full Insert Sequence. |
| | A_51_P461877 | Thbs2 | Thrombospondin 2 (Thbs2) |
| | A_52_P700056 | Wfdc17 | Wap Four-Disulfide Core Domain 17 (Wfdc17) |
| 53 genes: Stepwise downregulation | A_51_P337195 | Pipox | Pipecolic Acid Oxidase (Pipox) |
| | A_55_P2130338 | Olfr1135 | Olfactory Receptor 1135 (Olfr1135) |
| | A_51_P375783 | Prap1 | Proline-Rich Acidic Protein 1 (Prap1) |

| | | | |
|---|---|---|---|
| (miRs-normal ~ NC-normal ~ NC-polyp) | A_55_P1983162 | Pstpip2 | Proline-Serine-Threonine Phosphatase-Interacting Protein 2 (Pstpip2) |
| | A_52_P487436 | Nags | N-Acetylglutamate Synthase (Nags), Nuclear Gene Encoding Mitochondrial Protein |
| | A_55_P2042331 | Gzmk | Granzyme K |
| | A_52_P443334 | Cd8a | Cd8 Antigen, Alpha Chain (Cd8A), Transcript Variant 1 |
| | A_52_P638459 | Ccl5 | Chemokine (C-C Motif) Ligand 5 (Ccl5) |
| | A_51_P499195 | Nkg7 | Natural Killer Cell Group 7 Sequence (Nkg7) |
| | A_51_P487555 | Cd7 | Cd7 Antigen (Cd7) |
| | A_55_P2164443 | | Nap113319-1 |
| | A_51_P400366 | Rhbg | Rhesus Blood Group-Associated B Glycoprotein (Rhbg) |
| | A_52_P69558 | Gm8221 | Predicted Gene 8221 (Gm8221), Non-Coding Rna |
| | A_51_P498631 | Dfna5 | Deafness, Autosomal Dominant 5 (Human) (Dfna5) |
| | A_55_P1978502 | H2-Q1 | Histocompatibility 2, Q Region Locus 1 (H2-Q1) |
| | A_55_P1956160 | Gm8909 | Predicted Gene 8909 (Gm8909) |
| | A_55_P2107528 | | Chr8:107199491-107199432 |
| | A_55_P2293013 | Ces2a | Carboxylesterase 2A (Ces2A), Transcript Variant 1 |
| | A_55_P2101367 | Ppfia3 | Protein Tyrosine Phosphatase, Receptor Type, F Polypeptide (Ptprf), Interacting Protein (Liprin), Alpha 3 (Ppfia3) |
| | A_55_P1995339 | Btnl6 | Butyrophilin-Like 6 (Btnl6) |
| | A_55_P1996821 | | Myosin Xvb |
| | A_51_P243755 | Slc10a2 | Solute Carrier Family 10, Member 2 (Slc10A2) |

| | | | |
|---|---|---|---|
| | A_51_P499599 | Osr2 | Odd-Skipped Related 2 (Osr2) |
| | A_51_P283344 | 1700011H14Rik | Riken Cdna 1700011H14 Gene (1700011H14Rik) |
| | A_55_P1968848 | Gm4312 | Predicted Gene 4312 (Gm4312) |
| | A_55_P2069468 | Nt5c1b | 5'-Nucleotidase, Cytosolic Ib (Nt5C1B) |
| | A_52_P338066 | Ubd | Ubiquitin D (Ubd) |
| | A_55_P2073825 | Ocm | Oncomodulin (Ocm) |
| | **A_51_P405167** | **Maf** | **Avian Musculoaponeurotic Fibrosarcoma (V-Maf) As42 Oncogene Homolog (Maf)** |
| | A_52_P326664 | Unc93a | Unc-93 Homolog A (C. Elegans) (Unc93A) |
| | A_55_P1968443 | Naaladl1 | N-Acetylated Alpha-Linked Acidic Dipeptidase-Like 1 (Naaladl1) |
| | A_52_P2710 | Cml5 | Camello-Like 5 (Cml5) |
| | A_52_P669128 | Slc5a1 | Solute Carrier Family 5 (Sodium/Glucose Cotransporter), Member 1 (Slc5A1) |
| | A_51_P331288 | Akr1b7 | Aldo-Keto Reductase Family 1, Member B7 (Akr1B7) |
| | A_51_P502150 | Slc9a3r1 | Solute Carrier Family 9 (Sodium/Hydrogen Exchanger), Member 3 Regulator 1 (Slc9A3R1) |
| | A_55_P2115220 | Gm11437 | Predicted Gene 11437 (Gm11437) |
| | A_55_P2051778 | Sult2b1 | Sulfotransferase Family, Cytosolic, 2B, Member 1 (Sult2B1) |
| | A_55_P1974740 | Slc13a1 | Solute Carrier Family 13 (Sodium/Sulfate Symporters), Member 1 (Slc13A1) |
| | A_55_P1953583 | Ace2 | Angiotensin I Converting Enzyme (Peptidyl-Dipeptidase A) 2 (Ace2), Transcript Variant 1 |
| | A_55_P1988368 | Upp1 | Uridine Phosphorylase 1 (Upp1), Transcript Variant 1 |

| | | |
|---|---|---|
| A_55_P1965079 | | Predicted Gene 10499 |
| A_51_P161830 | Enpep | Glutamyl Aminopeptidase (Enpep) |
| A_51_P161265 | Ccl25 | Chemokine (C-C Motif) Ligand 25 (Ccl25), Transcript Variant 1 |
| A_51_P328631 | Sis | Sucrase Isomaltase (Alpha-Glucosidase) (Sis) |
| A_55_P2033795 | Fabp6 | Fatty Acid Binding Protein 6, Ileal (Gastrotropin) (Fabp6) |
| A_55_P1988488 | Npc1l1 | Npc1-Like 1 (Npc1L1) |
| A_55_P2117155 | Apoc3 | Apolipoprotein C-Iii (Apoc3) |
| A_55_P2181752 | Apol7a | Apolipoprotein L 7A |
| A_55_P2118436 | H2-T18 | Predicted: Histocompatibility 2, T Region Locus 18 (H2-T18) |
| A_55_P1962723 | | Nap022599-001 |
| A_55_P2162880 | Cyp3a57 | Cytochrome P450, Family 3, Subfamily A, Polypeptide 57 (Cyp3A57) |
| A_55_P2164102 | | Mrna For Immunoglobulin H-Chain V-Region, Partial Cds, Clone: 0S_A-12. |
| A_51_P240986 | Plekhg6 | Pleckstrin Homology Domain Containing, Family G (With Rhogef Domain) Member 6 (Plekhg6) |

MAF is shown in bold.

**Supplementary Table. S3. Relationship between c-MAF expression and clinical and pathological parameters.**

| | Strong (n = 35) | Weak (n = 63) | *P* value |
|---|---|---|---|
| Age (years) | | | |
| ≥ 65/< 65 | 17/18 | 35/28 | 0.507 |
| Gender | | | |
| Male/Female | 20/15 | 37/26 | 0.878 |
| Location | | | |
| Rectum/Colon | 11/24 | 32/31 | 0.064 |
| Depth | | | |
| T4/Tis, T1, T2, T3[a] | 2/33 | 7/56 | 0.375 |
| Lymph node metastasis | | | |
| Negative/Positive | 27/8 | 38/25 | 0.091 |
| Histological type | | | |
| tub1, tub2/muc, por[b] | 33/2 | 57/6 | 0.509 |
| Lymphatic duct invasion | | | |
| Negative/Positive | 17/18 | 12/51 | 0.002 |
| Venous invasion | | | |
| Negative/Positive | 27/8 | 39/24 | 0.123 |

a) Tis, carcinoma in situ; T1, involvement of submucosa; T2, involvement of muscularis propria; T3, involvement of subserosa; T4, involvement of serosal surface or direct invasion to other organs; b) tub1, well differentiated adenocarcinoma; tub2, moderately differentiated adenocarcinoma; muc, mucinous carcinoma; por, poorly differentiated adenocarcinoma

**Supplementary Table. S4. Lists of upregulation or stepwise downregulation genes between miRs-normal, NC-normal, and NC-polyp of *CPC;Apc* mice.**

| Upstream Regulator | Molecule Type | Predicted Activation State | Activation z-score | *P* value |
|---|---|---|---|---|
| **Vegf** | **Growth factor** | **Activated** | **5.108** | **< 0.001** |
| **IL4** | **Cytokine** | **Activated** | **4.659** | **< 0.001** |
| **IGF1** | **Growth factor** | **Activated** | **4.581** | **< 0.001** |
| **HGF** | **Growth factor** | **Activated** | **4.57** | **< 0.001** |
| **AGT** | **Growth factor** | **Activated** | **4.527** | **< 0.001** |
| **TRIM24** | **Transcription regulator** | **Activated** | **4.484** | **< 0.001** |
| RNASEH2B | Other | Activated | 4.413 | < 0.001 |
| **JUN** | **Transcription regulator** | **Activated** | **4.324** | **< 0.001** |
| **TGFB1** | **Growth factor** | **Activated** | **4.308** | **< 0.001** |
| **EGF** | **Growth factor** | **Activated** | **4.248** | **< 0.001** |
| **Insulin** | **Hormone** | **Activated** | **4.232** | **< 0.001** |
| SP1 | Transcription regulator | Activated | 4.102 | < 0.001 |
| **NRG1** | **Growth factor** | **Activated** | **4.041** | **0.001** |
| Ttc39aos1 | Other | Activated | 4.024 | < 0.001 |
| **STAT3** | **Transcription regulator** | **Activated** | **4.021** | **< 0.001** |
| **CITED2** | **Transcription regulator** | **Activated** | **4.02** | **< 0.001** |
| **FGF2** | **Growth factor** | **Activated** | **4.004** | **< 0.001** |
| NPM1 | Transcription regulator | Activated | 4 | < 0.001 |
| STAG2 | Other | Activated | 3.983 | < 0.001 |
| **NKX2-3** | **Transcription regulator** | **Activated** | **3.977** | **< 0.001** |
| PIK3CG | Kinase | Activated | 3.965 | < 0.001 |

Cancer-promoting growth factors, transcription factors, kinases, and cytokines are shown in bold.

**Supplementary Table. S5. Detail of nucleotides for qRT-PCR or generating *c-MAF* knock out mice**

| Primer sequence (5' to 3') or TaqMan Gene Expression Assay ID | | | |
|---|---|---|---|
| Mus musculus | c-MAF | | Mm02581355_s1 |
| | Actb | | Mm02619580_g1 |
| Rattus norvegicus | c-MAF | | Rn00824591_s1 |
| | Actb | | Rn00667869_m1 |
| Homo sapiens | c-MAF | | Hs04185012_s1 |
| | miR-155 | | 002623 |
| | RNU6B | | 001093 |
| | ACTB | | Hs01060665_g1 |
| | TP53 | F | GTTCCGAGAGCTGAATGAGG |
| | | R | TCTGAGTCAGGCCCTTCTGT |
| | CDKN1A | F | AAGACCATGTGGACCTGT |
| | | R | GGTAGAAATCTGTCATGCTG |
| | GATA3 | F | GCGGGCTCTATCACAAAATG |
| | | R | TCTGACAGTTCGCACAGGAC |
| | ACTB | F | GATGAGATTGGCATGGCTTT |
| | | R | CACCTTCACCGTTCCAGTTT |
| List of oligonucleotide sequence (5' to 3') for CRISPR Cas9 system or genotyping | | | |
| gRNA | | | CAGGAGGATGGCTTCAGAAC |
| PCR primer | | F | GTGTGCACGTTCGAGCTTT |
| | | R | GTCATCCAGTAGTAGTCTTCCAGGT |
| Sanger sequence primer | | R | CAGGTGCGCCTTCTGTTC |
